# Mapping the Complex Genetic Landscape of Human Neurons

**DOI:** 10.1101/2023.03.07.531594

**Authors:** Chen Sun, Kunal Kathuria, Sarah B Emery, ByungJun Kim, Ian E. Burbulis, Joo Heon Shin, Brain Somatic Mosaicism Network, Daniel R. Weinberger, John V. Moran, Jeffrey M. Kidd, Ryan E. Mills, Michael J. McConnell

## Abstract

When somatic cells acquire complex karyotypes, they are removed by the immune system. Mutant somatic cells that evade immune surveillance can lead to cancer. Neurons with complex karyotypes arise during neurotypical brain development, but neurons are almost never the origin of brain cancers. Instead, somatic mutations in neurons can bring about neurodevelopmental disorders, and contribute to the polygenic landscape of neuropsychiatric and neurodegenerative disease. A subset of human neurons harbors idiosyncratic copy number variants (CNVs, “CNV neurons”), but previous analyses of CNV neurons have been limited by relatively small sample sizes. Here, we developed an allele-based validation approach, SCOVAL, to corroborate or reject read-depth based CNV calls in single human neurons. We applied this approach to 2,125 frontal cortical neurons from a neurotypical human brain. This approach identified 226 CNV neurons, as well as a class of CNV neurons with complex karyotypes containing whole or substantial losses on multiple chromosomes. Moreover, we found that CNV location appears to be nonrandom. Recurrent regions of neuronal genome rearrangement contained fewer, but longer, genes.

## Introduction

It is inaccurate to view an individual’s genome as invariant from organ to organ, or from cell to cell within an organ. For example, somatic mosaicism among lymphocytes has been recognized since the 1970’s with the discovery of somatic gene rearrangement at T cell receptor and immunoglobulin loci ^1^. Recurrent somatic mutations also underlie the pathology of many cancers ^2^. Recent advances in single cell and bulk DNA sequencing approaches have revealed abundant somatic mosaicism throughout the human body^3, 4, 5, 6, 7, 8^. Associated studies have linked environmental mutagens to somatic mutations in the skin, bladder, and other exposed cells ^6, 9, 10^. Rapidly dividing stem cell populations also incur somatic mutations due to DNA replication errors. Clonal expansion of variant genomes can, in turn, shape mosaicism among an individual’s somatic cells ^11^. Somatic mutations, accompanied by cell death, set the stage for somatic selection during the lifespan of an individual.

Brain somatic mosaicism is associated with neurodevelopmental disorders, especially epilepsy ^12, 13, 14, 15, 16, 17, 18, 19^. Unlike other organs, cerebral cortical neurons arise *in utero* and are not replaced during normal human lifespan ^20^. Neural stem and progenitor cells proliferate rapidly during human cortical development; these progeny overpopulate the developing cerebral cortex. Somatic selection is one means by which some progeny may thrive as cortical neurons while other progeny perish ^21, 22, 23, 24^. The genomes of mature cortical neurons contain hundreds of single nucleotide variants (SNVs), some of which mark clonal lineages ^25, 26, 27, 28^. LINE-1 mobile elements retrotranspose during neurogenesis and contribute to brain somatic mosaicism in a small subset of neurons ^29, 30, 31, 32, 33^. Although SNVs are numerous and accumulate throughout life, relatively few are predicted to have protein-coding mutations with obvious consequences for affected neurons^28, 34, 35^. Megabase (Mb)-scale copy number variants (CNVs) - typically sub-chromosomal deletions - also contribute to brain somatic mosaicism ^36, 37, 38^.

In non-diseased (neurotypical) brains, dozens of genes are impacted in CNV neurons with substantial inter-individual variability in the frequency of CNV neurons among individuals ^39^. CNV neurons are more prevalent in the frontal cortex of young individuals (n=4 individuals <30 years old; 28.5% CNV neurons, 75/263) than in aged individuals (n=5 individuals >70 years old; 7.3% CNV neurons, 26/354) ^39^. However, small sample sizes (<100 neurons / individual) have limited the ability of these studies to find patterns of recurrent rearrangement (*e*.*g*., CNV hotspots) among neuronal genomes. If present, such recurrent sites of neuronal genome rearrangement may provide insight into the mechanisms and/or consequences of brain somatic mosaicism. Recurrent sites of neuronal genome rearrangement could be influenced by common fragile sites that are predisposed to genome rearrangements ^40, 41^ and may reflect neurodevelopmental somatic selection. Neither mechanism precludes the other.

We reasoned that if recurrent brain CNVs exist, hotspots would be found among neurons in any millimeter-scale cortical biopsy from a single individual. Using a commercial droplet-based whole genome amplification (WGA) method, we generated Illumina sequencing libraries from 2,125 frontal cortical nuclei isolated from a previously characterized neurotypical individual ^28, 39^. Read-depth analysis of each cell was coupled with phased germline single nucleotide polymorphisms (SNPs) to develop a single cell sequencing coverage and allele-based approach (SCOVAL) that restricted read-depth based deletion calls using concordant, phased, loss-of-heterozygosity (LOH) information. In total, 2097 single neuron libraries passed quality controls (QC) and 10.8% (226/2097) contained at least one Mb-scale CNV. An unexpected subpopulation of these CNV neurons (65/226, 25%) have highly aberrant karyotypes wherein multiple chromosomes harbor multiple deletions, including 6 aneusomic neurons. When compared to a random model, CNVs are depleted in gene-dense genomic regions. However, frequent neuronal genome rearrangements are more common in genomic regions that contain genes encoded by more than 100 kilobases (kb) of genomic sequence (herein defined as long genes).

## Results

### Determining the genetic architecture of individual neurons

When CNVs are clonal or recurrent, as in populations of cancer cells, read-depth based single cell genomic approaches can accurately reconstruct clonal cell lineages ^42, 43, 44^.

However, neuronal CNVs are rarely clonal ^39^ precluding validation in lineage-derived “sister” neurons. SCOVAL combined read-depth and phased LOH metrics (**Fig. 1A)** to determine the prevalence of CNVs in single neurons isolated from the post-mortem brain of a neurotypical 49-year-old male. Samples from the same neurotypical individual were analyzed in previous studies conducted by the Brain Somatic Mosaicism Network ^28, 45^ and a directly relevant small study (*i*.*e*., consisting of 99 neuronal nuclei, 26 non-neuronal nuclei) that identified 11 CNV neurons (∼11%) and two non-neuronal nuclei (7.6%) containing CNVs > 2Mb ^39^.

**Figure 1.**
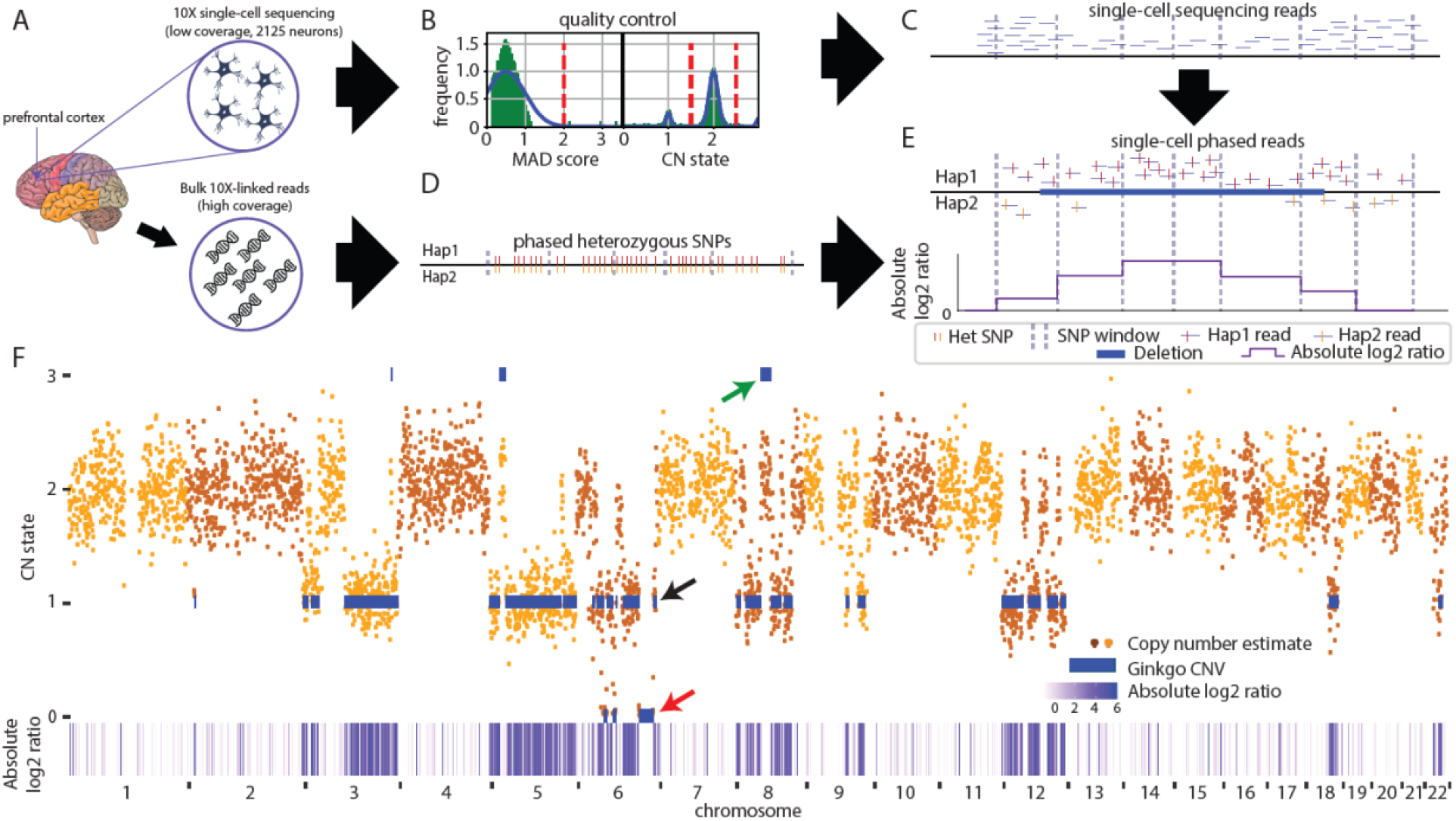
SCOVAL: identification of copy number variation using read-depth and allele imbalance. Overview of SCOVAL. (**A**) Single nuclei and bulk dural fibroblast DNA were analyzed using 10X platforms. (**B**) Single nuclei library quality is assessed based on median absolute deviation (MAD) and copy number thresholds are established using population statistics. Graphs depict schematized data; vertical red lines illustrate threshold strategy. (**C**) Candidate CNVs are identified based on altered read depth across consecutive genomic bins. Heterozygous SNPs are phased using bulk linked-reads in chromosomal segments (“hap 1” or “hap 2”). (**E**) Absolute log2 ratios derived from “hap1” / “hap 2” are calculated across ∼100 SNP windows (see text). A deletion with concordant loss of heterozygosity (log2 ratio <> 0) is illustrated. **(F)** A highly aberrant CNV neuron (#5) shows representative Gingko calls (blue bars), duplications (*e*.*g*., green arrow), heterozygous deletions (*e*.*g*., black arrow), and homozygous deletions (*e*.*g*., orange arrow) and qualitatively concordant increases in absolute log2 ratio (white<purple). The genome is plotted from left to right on the x axis, read-depth is in the upper panel (CN state on the Y axis) and absolute log2 ratios are reported in the lower panel.

Briefly, we isolated >50,000 human frontal cortical neurons using fluorescence-activated nuclei sorting of NeuN-positive nuclei. Two DNA libraries were then prepared in separate lanes on the 10X Genomics Chromium platform (**Fig. 1A**); each lane obtained ∼1,000 single neuronal genomic libraries with unique barcodes. The resultant libraries (2125 total) were combined into one pool that was sequenced in two batches on an Illumina NovaSeq platform, achieving an average of 2.83 +/-1.22 million reads per neuron. Following our previous approach ^39^, we mapped reads to 5067 variable sized autosomal bins, each containing 500kb of uniquely mappable sequence (mean bin size = 569kb, range = 501 to 2812kb). Our quality control (QC) filters excluded 28 single neurons with aberrant bin-to-bin variance [*i*.*e*., Median Absolute Deviation (MAD), 2097 (>95%) libraries passed QC] and masked 308 genomic bins that were outliers in global read coverage across all neurons (**Supplementary Fig. 1A-C**). We adapted Ginkgo ^44^ to call CNVs larger than 1Mb, defined copy number (CN) state thresholds (**see Methods**), and identified 2,564 putative autosomal CNVs (2,401 deletions and 163 duplications) in 469 different neurons **(Fig.1B, Supplementary Table 1**).

In parallel, we sequenced dural fibroblast DNA from the same individual at high coverage (∼52.7X) to identify and phase germline SNPs using 10X Genomics linked-read sequencing ^46^. Briefly, this approach isolated and fragmented long DNA segments into barcoded short reads that could be used to reconstruct underlying haplotypes into 2548 phased genomic blocks (mean 1178kb +/-2034kb, median 234kb). Within each of these phased blocks, we further segmented the genome into windows of 20-100 phased heterozygous germline SNPs (mean = 107kb, range = 0.687 to 1470kb) that arbitrate predicted somatic deletions with phased LOH. For each window of each cell, we counted the number of informative reads (*e*.*g*., reads that intersect with phased heterozygous SNPs) on each haplotype. We then calculated the absolute log2 ratio of the number of reads on each haplotype and integrated this ratio into the filtering models (**Fig. 1C**). The application of our naïve Bayesian-based pipeline (**see Methods, Supplementary Fig. 2**) identified 1,985 regions with both sequence coverage and phased LOH support consistent with heterozygous deletions in 231 neurons. We excluded Gingko deletion calls where more than 75% of internal phased SNP windows contained fewer than 3 informative reads and arrived at a call set of 1,853 heterozygous somatic deletions in 226 neurons.

Other candidate neuronal CNVs (*i*.*e*., duplications and homozygous deletions) were more challenging to validate using SCOVAL. Previous studies using read-depth alone reported more than two-fold fewer duplications than deletions ^36, 39^. Using SCOVAL, we measured allelic ratios between haplotypes to assess the 163 Ginkgo duplication calls. The log2 ratios of haplotype-resolved alleles for each duplication were not significantly different from randomly sampled euploid regions of that particular cell (one-tailed t-test, p-value = 0.998, **Supplementary Fig. 3A**). These findings suggest that greater single cell sequencing coverage may be required for SCOVAL to assess duplications in single neuron WGA data, although phased LOH may also allow us to filter regions where Ginkgo reports false positives (**Fig. 1F**, green arrow). Nevertheless, although some of these regions may represent *bona fide* duplications, set we opted to exclude putative duplications with only Gingko support from further analysis in the interest of evaluating a conservative call.

Homozygous deletions have been uncommon in previous datasets and have distinct properties compared to heterozygous deletions. Specifically, these deletions are not directly amenable to allelic modeling as both haplotypes are absent and any observed non-zero allele ratios likely would be derived from mis-mapped reads. Thus, we developed an additional filter to reduce the false positive rate for 106 putative homozygous deletions with read-depth support.

We calculated a read-depth ratio for each Ginkgo window by comparing the read-depth in every cell with the read-depth from bulk sequencing ^28^ and derived a Gaussian mixture model to calculate the posterior probability for putative homozygous deletions using these values from our initial heterozygous and homozygous deletion calls (see **Methods, Supplementary Fig. 3B**) This strategy found additional support for 86/106 putative homozygous deletions (posterior probability > 0.99, **Supplementary Fig. 3C**). These 86 regions were included in our final deletion call set for subsequent analyses of CNV locations. Importantly, homozygous deletions are only found in neurons with highly aberrant karyotypes and all flank a heterozygous deletion (**Fig. 1F**, red arrow), indicating that they are likely the result of two independent and overlapping heterozygous deletions. Further, we identified 8 Ginkgo-called homozygous deletions that exhibited a read depth and allele ratio profile consistent with heterozygous deletions and reclassified them as such (**Supplementary Fig. 4**).

SCOVAL produced a final deletion CNV set (**Supplementary Table 2)** comprising 1,957 somatic CNV calls (13.95 Mb +/-17.47 Mb) among 226 CNV neurons (∼11%). These represent 76.3% of the initial 2,564 read depth predictions. CNV neuron prevalence (226/2097 neurons) is in good agreement with previous read-depth based CNV detection from this individual (∼11% of 99 neurons) ^39^. Although the nature of single-cell DNA sequencing prohibits the direct validation of identified CNVs, manual, subjective inspection of read-depth and allele ratios are strikingly concordant.

SCOVAL was designed to identify idiosyncratic CNVs in human neurons. Another single-cell CNV caller, CHISEL, was designed to study tumor evolution and intra-tumor heterogeneity ^47^. CHISEL and similar approaches ^48^ assume a higher frequency of tumor subclones (>5-10% ^49^) than has been observed in CNV neurons ^39^. When we tested CHISEL using our single neuron data, almost all reported CNVs (21,906) clustered collectively within 12 genomic loci (99.25% of CHISEL calls) and were reported in more than 50% of neurons (**Supplementary Fig. 5**). Notably, 11 of the 12 loci overlapped with SCOVAL outlier bins that were associated with WGA artifacts (**see Methods** and ^39^). We compared the remaining 165/21906 CHISEL CNV calls with our final call set. These 165 calls were reported in only three neurons, but 39 CHISEL CNV calls overlapped with 15 SCOVAL CNV calls. Manual inspection of read-depth and LOH at the other 126 CHISEL CNV calls found no subjective support (**Supplementary Table 3)**. Consistent with reports attempting to apply similar cancer-oriented approaches for identifying somatic CNVs in neurons ^28^, we conclude that CHISEL and other cancer-oriented approaches are not appropriate to study brain somatic mosaicism.

### Some CNV Neurons have highly aberrant karyotypes

SCOVAL identified 226 CNV neurons with at least one deletion. These deletions ranged in size from 1Mb to entire chromosomes. We also observed that when neurons harbored multiple deletions, many clustered on single chromosomes. In contrast to a uniform background model (see **Methods** and below), CNVs did not appear to be distributed randomly among CNV neurons (**Fig. 2A**). Forty-six CNV neurons contained a single deletion, but five contained greater than 30 deletions. Apparent chromosomal monosomies (*i*.*e*., where all genomic bins reported a copy number (CN) state = 1) were observed in six different neurons. One neuron (#1) was monosomic for Chr5, another (neuron #7) was monosomic for Chr9, two neurons (#2, 3) were monosomic for Chr13, and two other neurons (#4, 46) were monosomic for Chr18 (**Fig. 2B, C**). All monosomic neuronal genomes were highly aberrant and harbored many additional deletions affecting 40 – 98% of other chromosomes (**Fig. 2C**). Among 65 CNV neurons with deletions affecting >5% of their genome, 48 contained at least one chromosome that was >50% monosomic.

**Figure 2.**
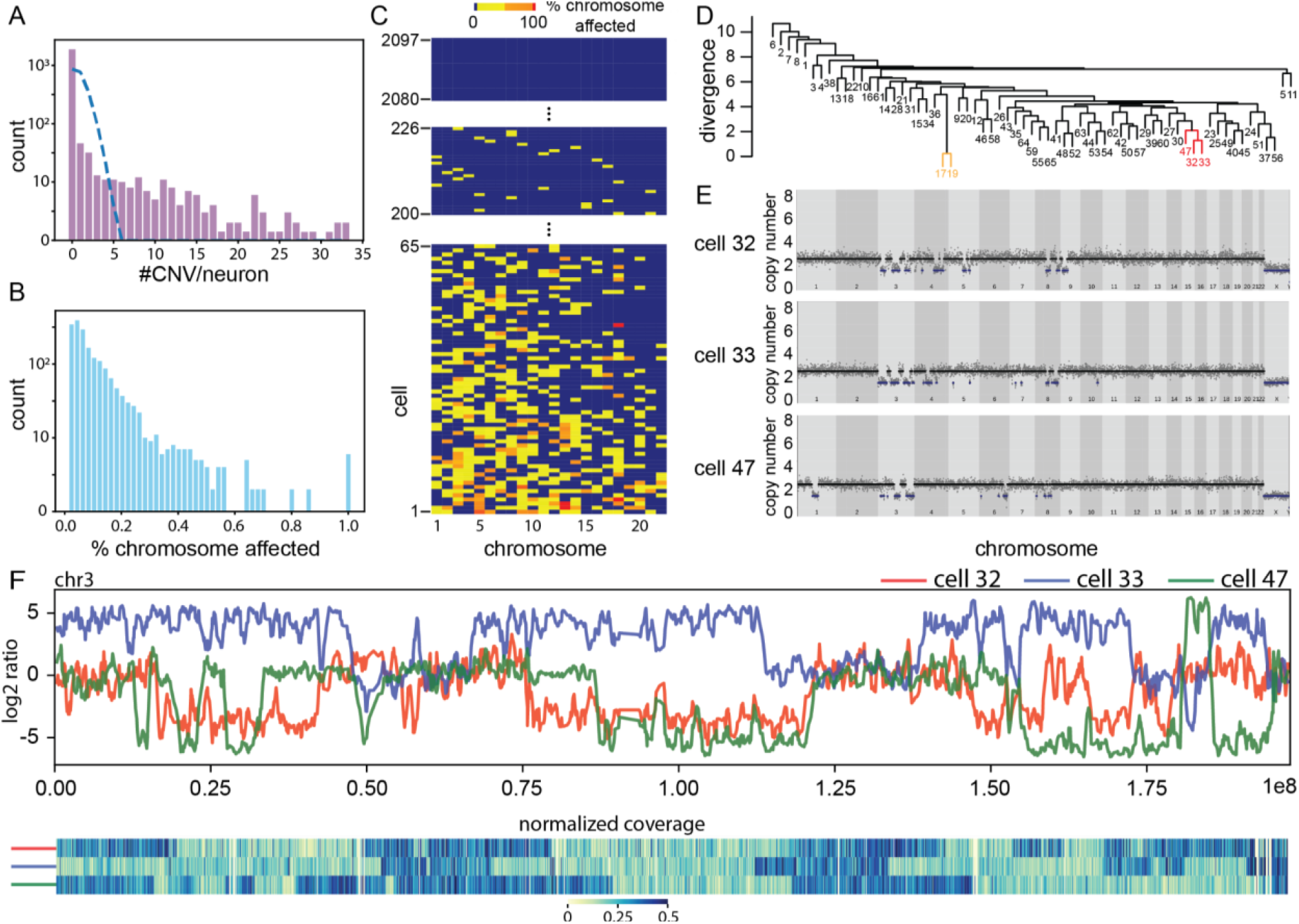
CNV neurons can have highly aberrant karyotypes. **(A)** The observed CNV per neuron [(purple bars, counts (y axis),CNVs/neuron (x axis)] distribution deviates (P < 0.0001) from Poisson expectations (dashed blue line). **(B, C)** Deletions cluster in a subset of CNV neurons. (B) Counts (y axis) of the cumulative percent of each chromosome deleted (n = 2097 neurons * 22 autosomes) in CNV neurons. (C) Neuronal genomes (n=2097) are arranged in a cells-by-chromosome matrix, ranked by the total percentage of their genome containing deletions. Cell #226 is the first CNV neuron among 2097 total neurons with the smallest observed single deletion (blue = unaffected chromosome, yellow <50%, orange = 50 - 99%, red 100%). **(D-F)** Among 65 neurons with the most aberrant genomes, some have similar karyotypes. (D) Hierarchical clustering identifies two groups (yellow, red) with the least divergence from similarity (y axis). (E) Red cluster neurons [cells #32, 33, and 47 in (C)] have similar CNV profiles. Read-depth is plotted as in Fig. 1F. The yellow cluster (cells #17 and #19) is shown in Fig. S6B. (F) Concordant read-depth is observed on opposite haplotypes in the most similar pair [#32(red) and #33(blue)]. When overlapping, events on cell #47 (green) match the #32 haplotype, but never the #33 haplotype. Chromosome 3 is plotted from left to right. Haplotype log2 ratio (upper panel) and corresponding read-depth (lower panel, blue = diploid) plots show overlapping deletions and LOH for each haplotype.

**Figure 3.**
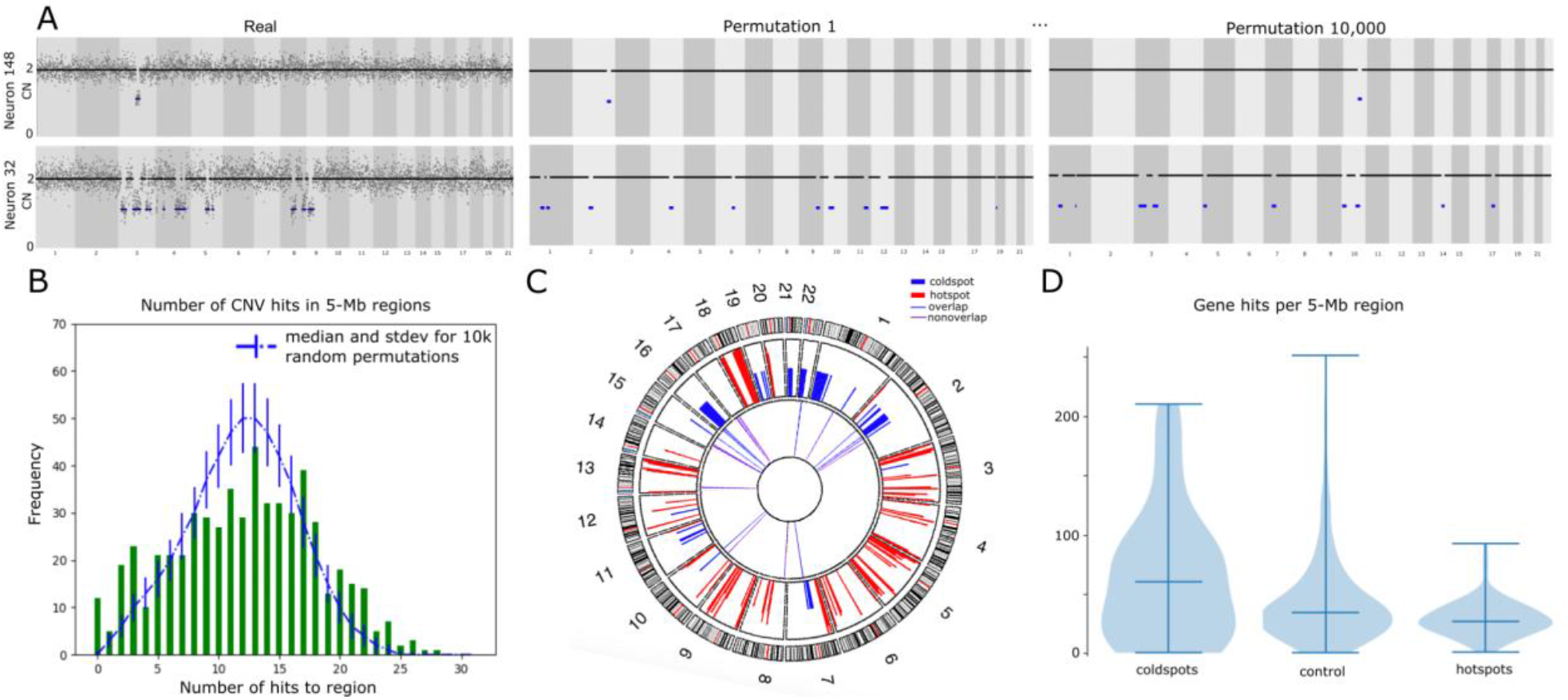
Analysis of CNV distribution relative to random null model. **(A)** Empirical read-depth plots of two CNV neurons (left panels) and representative permutations (right two panels) are displayed as in Fig. 1F. **(B)** Relative to 10,000 permutations of real data (represented by blue dotted line and error bars), high and low CNV burden are enriched at the extremities of the Gaussian distribution (green bars). **(C)** Circos plot shows that hotspots (red, outer tier) and cold spots (blue, middle tier) cluster on distinct chromosomes. Thirty-three pathogenic CNVs (blue, purple, inner tier) never overlap hotspots. Eleven (blue) overlap cold spots. **(D)** Violin plot showing gene enrichment in cold spots (left) and depletion in hotspots (right) relative to other 5Mb regions (P<0.001).

We evaluated CNV locations in CNV neurons based on the percentage of each chromosome affected by CNVs (**Fig. 2C**) and found two pairs of neurons (#17, #19 and #154, #155) that were nearly identical in their genomic read-depth patterns and could, in principle, represent clonal “sister” neurons that arose from a common progenitor cell during neurodevelopment (**Supplementary Fig. 6**). However, each of these pairs arose from the same 10X Genomics Chromium lane; therefore, we cannot exclude the possibility that one nucleus may have paired with two 10X GEM beads in a single droplet. Subsequent analyses assume that these two pairs are highly concordant technical replicates.

Hierarchical clustering (**Fig. 2D**) identified three other neurons (cells #32, #33, and #47) with similar karyotypes that could, in principle, share identity by descent (**Fig. 2E**). Thus, we investigated whether these deletions occurred on the same chromosomal phase block *(i*.*e*., haplotype). Multiple deletions in cells #32, #33, and #47 mapped to Chr3; however, read-depth alone cannot assess whether these deletions occur on the same physical chromosome. Also, 10X linked-read haplotyping identifies phased SNPs with Mb-scale resolution, as described above. To determine phasing at a chromosome level, we generated extended phase blocks using three CNV neurons (cells #33, #10, and #5) that contained overlapping deletions accounting for the full-length of Chr3 (**Supplementary Fig. 7**). Although CNV locations overlapped among these three neurons (**Fig. 2F**), the Chr3 CNVs were constrained to one haplotype in two neurons (cells #32 and #47) but occurred on the other haplotype in the third neuron (cell #33). The presence of other idiosyncratic CNVs suggest that these three neurons arose in distinct neurodevelopmental lineages. The possible ontogeny of these chromosomes might include chromosome mis-segregation, micronucleus formation, and a chromothripsis-like event ^50, 51, 52, 53, 54^. In any case, the strikingly similar patterns of loss observed in these three neurons likely represent recurrent rather than clonal events.

### CNVs are not randomly distributed in neuronal genomes

The similar patterns of chromosomal loss observed in subsets of CNV neurons led us to hypothesize that, in contrast to what has been reported in other tissue types ^55^, neuronal CNV locations may not arise randomly. Thus, we generated a control dataset of randomly placed deletions and explored whether neuronal genomes accumulate CNVs in “hotspots” or are protected from CNVs in “cold spots.” Briefly, the empirical call set was randomly rearranged, without collision, while keeping the size and abundance of CNVs constant on a per neuron basis. We reasoned that randomly, and reiteratively, placing the “real” CNVs throughout the genome would effectively generate a “random” CNV landscape (**Fig. 3A**); we then performed 10,000 synthetic iterations of real data to generate a null model. For analysis, the genome was segmented into 567 contiguous 5Mb regions and the number of simulated CNVs that overlapped each 5Mb genomic region (*i*.*e*., hits) were counted to generate a null model.

A Gaussian-shaped distribution of CNVs / 5Mb region was observed in the null model, but empirical data was enriched for observations at the extremities (**Fig. 3B**). Specifically, when empirical *P*-values were calculated for each 5Mb region, we found eighty-three 5Mb regions (14.6%) where observed CNVs occurred more frequently than in the random model (“hotspots,” *P*-value <0.05) and fifty-six 5Mb regions (9.9%) where empirical CNVs overlapped less frequently than in the null model (“cold spots,” *P*-value >0.99) (**see Methods for *P*-value determinations**). For example, fourteen 5Mb regions were hit at least 24 times by real CNVs, however this frequency (≥24 hits in a 5Mb region) occurred in only 0.5% of null model permutations. Importantly, no CNV-free region was observed in null model perturbations, but seven CNV-free cold spots were found in empirical data.

CNV hotspots and cold spots also clustered in several semi-contiguous stretches of the genome (**Fig. 3C**). Eighty-three 5Mb hotspots clustered into 47 distinct contiguous regions, whereas the 56 cold spots clustered into 22 distinct contiguous regions. Surprisingly, individual chromosomes also clustered as either hot or cold with respect to CNV presence or absence.

For example, 9/83 (∼11%) and 15/83 hotspots (∼18%) clustered on chromosomes 18 and 5, respectively, whereas 12/56 cold spot regions (21%) clustered on chromosome 1. Thirteen highly aberrant neuronal genomes (containing ≥25 CNVs in empirical data) all had a CNV(s) that intersected hotspots, whereas only nine had CNVs intersecting cold spots. Similarly, of the 112 CNV neurons that contained between 1-5 CNVs, fifty-four had CNVs intersecting hotspots and only seven had CNVs intersecting cold spots. Overall, 163 neuronal genomes had a CNV(s) overlapping a hotspot, whereas only 50 CNV neurons overlapped cold spots.

Because a depletion of CNVs in some regions could artificially increase the detection of CNVs elsewhere, putative CNV cold spots and hotspots may have a technical explanation.

Thus, we also functionally assessed observed cold spots for overlap with 33 germline CNVs (fifty-six 5Mb regions) that are associated with adverse neurodevelopmental phenotypes ^56^. One third (11/33) of these germline CNVs were in cold spots. By comparison, none (0/33) of the germline CNVs overlapped hotspots. The probability that a neuropathogenic germline CNV occurs in any 5Mb genomic region by chance is approximately 33/567 (5.8%); however, empirical overlap was observed in 11/56 (19.6%) 5Mb cold spot regions. Gene content further distinguished hotspots and cold spots from other control regions of the genome (**Fig. 3D**). Cold spots typically were gene dense (64.7 +/-56.2 genes per 5Mb region) and were not distributed uniformly when compared to control regions of the genome. By comparison, hotspots typically were gene-sparse relative to cold spots (32.6 +/-15.2 genes per 5Mb region).

### Recurrent regions of neuronal genome rearrangement

The observation that neuronal deletions cluster in genomic hotspots suggested that local genomic instability could, in principle, lead to recurrent mosaicism among neurons. To explore this hypothesis, we examined CNV start or end locations (*i*.*e*., breakpoints) that were shared amongst CNV neurons. Breakpoints are defined by one of the 5067 variably sized Gingko bins that each include 500kb of mappable sequence. Among these bins, 857 accounted for two or more CNV breakpoints (termed CNVBs) (**Fig. 4A and 4B)**, many of which (220/851; ∼26%) fell within previously identified hotspots.

**Figure 4.**
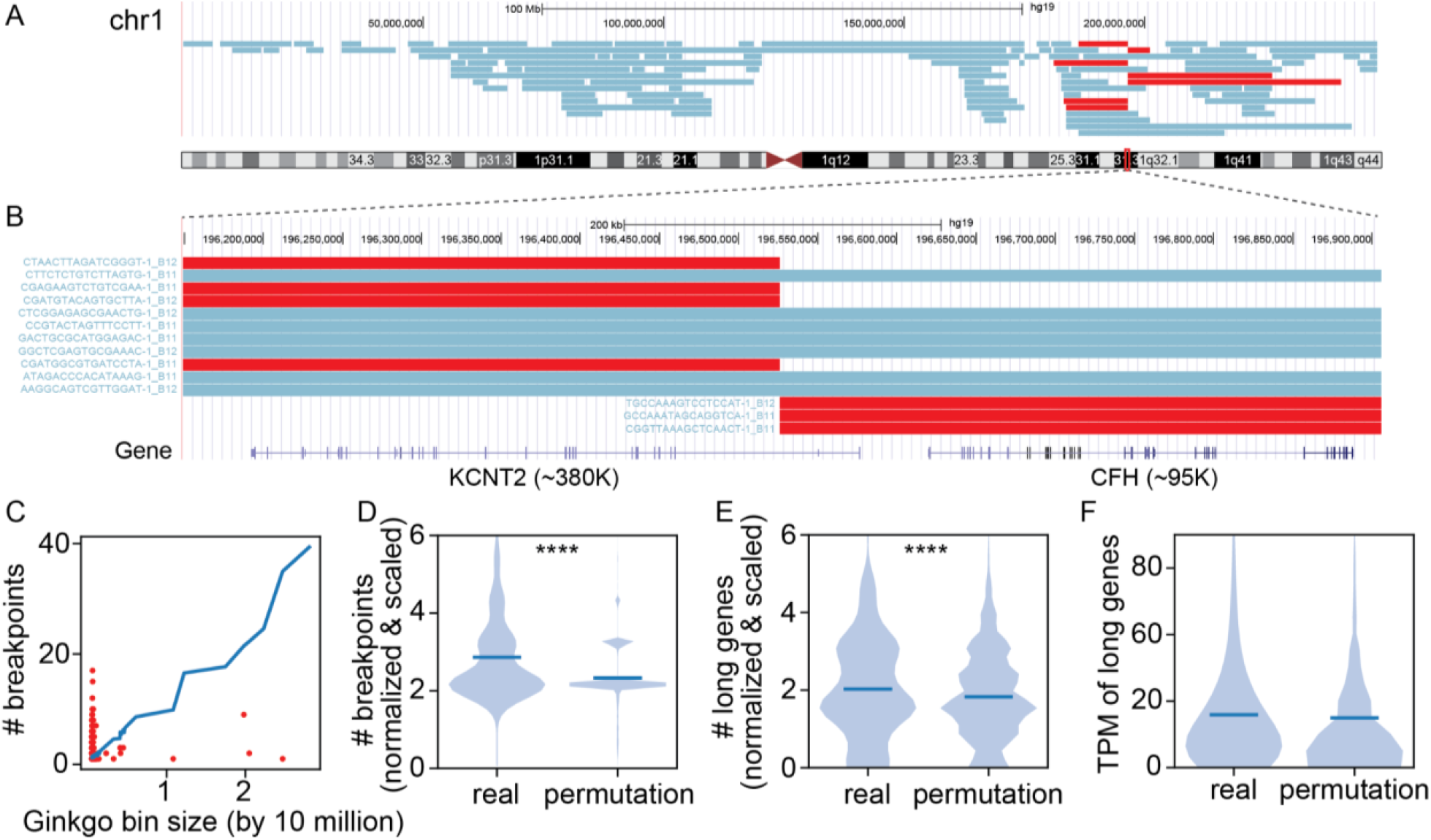
Recurrent CNV breakpoints across multiple neurons. **(A)** UCSC Genome Browser view of all CNVs detected on Chromosome 1 (47 neurons, rows). Seven neurons (red) contain CNVs that share a breakpoint region (CNVB). **(B)** Representative CNVB (red) on Chromosome 1 overlaps (+/- 250kb) two genes (lower panel). **(C)** Number of breakpoints identified in each Ginkgo bin (y axis) relative to bin size (x axis), shown for bins containing two or more CNVs (red) and averaged across all permutations in control set (blue line) **(D-F)** Violin plots show real and permuted data sets, normalized by bin size, when examined for (D) number of breakpoints, (E) number of long (>100k) genes (**** p<0.0001 for one-tailed t-test), and (F) transcripts per million bp (TPM) values of the longest gene in each bin.

We next sought to determine whether the number of bins containing more than 2 breakpoints was significantly different from a random CNV distribution (*i*.*e*., the control set of CNV permutations). Given variably sized Gingko bins (**Methods**), we first assessed whether Ginkgo bin size impacted breakpoint frequency. While bin size scaled linearly with CNVB frequency in random permutations, this linear relationship was not observed with empirical CNVBs (**Fig. 4C**). When breakpoint counts are normalized by bin size, observed CNVBs cluster more frequently in common bins than random CNVBs (one-sided t-test, *P*-value: 2.08*e-134), suggesting that CNVBs likely originate from a non-random process (**Fig. 4D**).

Empirical CNVBs were further assessed for properties that might suggest mechanisms of CNV formation. As the endpoints of each CNV are imprecise within the Ginkgo bins, we were unable to use typical approaches that examine sequence context around precise structural breakpoints ^57^; thus, we restricted our analysis to larger genomic features. Recent studies have indicated that somatic CNV hotspots in non-cancer systems are localized around large (>500kb) transcriptional units that form due to replication stress by a mechanism termed transcription-dependent double-fork failure ^58, 59^. To test if the CNVBs in our empirical dataset were consistent with this mechanism, we examined gene content in CNVB regions relative to random CNV permutations. Intriguingly, we observed a significant enrichment of empirical CNVBs within long genes (which we define as >100kb, one-sided t-test, *P*-value: 1.32*e-5), suggesting possible support for the hypothesis ^60, 61, 62^ that longer genes may incur an increased frequency of DNA double strand breaks (DSBs) and, in turn, lead to neuronal CNVs (**Fig. 4E**). However, the size of our detected CNVs and corresponding breakpoint windows are large and CNVB locations were not enriched for expression level (**Fig. 4F**). Thus, neuronal CNVs could arise by related, but perhaps different, mechanisms.

Among 98 of the 226 CNV neurons, we observed 73 CNVs that shared both 3’ and 5’ CNVBs. These may be recurrent CNVs (CNVRs). Haplotype information was then used to determine if CNVRs support a clonal relationship among neurons. Briefly, we used phased allele ratios to compare whether CNVRs shared haplotypes by determining the median of the differences between the minimum and maximum log2 allele ratios observed in each SNP window within the CNVR across all cells where it was identified, reasoning that lower log2 allele ratio values would represent CNVRs on a shared haplotype (**Methods, Supplementary Fig. 8A**). These calculations resulted in two apparent distributions of both lower (32/73) and higher (41/73) delta log2 ratio values. The lowest delta log2 ratio cluster contained the two pairs of technical replicates (**Fig. S5**), indicating the veracity of our approach. The remaining CNVRs exhibited a delta median log2 ratio larger than 5, suggesting that these CNVs occurred on opposite haplotypes (**Supplementary Fig. 8B**). However, all CNV neurons harboring CNVRs had complex karyotypes with divergent CNV patterns across the genome (*e*.*g*., **Fig. S13**).

These findings suggest that shared CNVs are not necessarily clonally-derived, but, instead, likely represent recurrent events (**Supplementary Fig. 8C**,**D**). Of note, similar CNVRs were observed in the analysis of cancer genomes and are referred to as “mirrored-subclonal” CNVs ^47, 63^.

## Discussion

The genetic landscape of human neurons is a mosaic of the individual’s germline genome; it is likely that every human neuron accumulates more than a thousand somatic variants over a person’s lifetime ^45, 64, 65, 66^. Specific somatic mutations have been linked to overgrowth phenotypes in patients with hemimegalencephaly and focal cortical dysplasia ^15, 67, 68,69^. Other studies report differential somatic mutation burden in subsets of patients with autism and schizophrenia ^12, 19, 70^. Furthermore, mosaic SNVs mark neural cell lineages within brain regions and some neuronal SNVs trace their origin prior to neuroectodermal specification ^4, 26, 28^. Somatic SNVs acquired in early human development can be shared among progeny in multiple germ layers. Mosaic SNVs contribute to intra-individual genetic variation, but mosaic Mb-scale CNVs alter the neurogenetic landscape in dramatic ways. However, it is unknown whether some genomic regions are more, or less, prone to CNV occurrence than other regions. The identification of CNV-prone genomic loci, if they exist, could indicate mechanisms for somatic CNV formation, and, possibly, reveal a role for CNV neurons in brain function and disease.

Here we employed a droplet-based WGA approach to map CNVs in 2097 frontal cortical neurons from a single individual. Technical barriers have limited previous studies to extrapolation from fewer than 100 neurons per individual and reported a total of 129 CNV neurons out of 879 frontal cortical neurons examined among 15 individuals^39^. We developed SCOVAL to add veracity to read-depth based CNV detection through an analysis of haplotype imbalance. We showed high concordance between heterozygous deletions identified by read-depth and by phased LOH in single neuronal nuclei. In this sample, we found that 226/2097 (10.8%) of neurons harbor at least one Mb-scale CNV, and that 2% of CNV neurons exhibited aneuploidies. Moreover, we found that 65/226 CNV neurons contained many deletions across multiple chromosomes leading to highly aberrant karyotypes.

By combining haplotype and read-depth approaches, we have strong confidence that neuronal genomes contain large segments of chromosomes that are not sampled using single cell sequencing approaches. This finding is consistent with previous reports that have examined a limited number of cells from neuronal and non-neuronal tissues using multiple technologies.

Although we posit that the assayed sequence is missing because the corresponding segments have been deleted *in vivo*, unexpected technical or biological factors may yet contribute to the loss of signal. For example, neuronal preps exclude micronuclei ^71^ however, the appreciable occurrence of micronuclei in neuronal tissue would still reflect an underlying alteration in genome content in the brain. Similarly, the lack of validated duplications in single cell neuronal sequencing is striking. Further study is required to develop mechanistic or technical explanations for this disparity.

Our finding of a nonrandom distribution of 1,861 deletions among 226 CNV neurons also allays concerns of random technical artifacts in neuronal CNV detection. Spurious WGA events, such as uneven genome amplification, are expected to occur randomly across the genome and are physically limited in size by the processivity of the polymerase (<20kb). Multiple whole genome amplification (WGA) approaches have been performed on single human neurons; all of these reported Mb-scale CNVs ^36, 37, 38^. This technical concern was addressed previously ^39, 72^ wherein a similar prevalence of CNV neurons was observed in two samples from the same individual (26 year-old), subjected to different WGA approaches. In Chronister, *et al*., parameter optimization on synthetic datasets limited read-depth based CNV detection to false positive rates <5%. Here, we provide additional lines of evidence that single cell approaches for neuronal CNV detection are robust to technical artifacts. First, we showed that SCOVAL finds haplotype allele-level support for 76% of read-depth based deletion calls. Importantly, 99% of >10 Mb heterozygous deletions received orthogonal support via phased LOH. Second, when SCOVAL was applied to 2,097 neurons, the fraction of CNV neurons observed (10.8%) was concordant with the fraction (11.1%) identified using different chemistry on a smaller (99 neuron) sample from the same brain region. Perhaps most strikingly, we identified CNV hotspots and cold spots that were inconsistent with a random distribution of technical artifacts. Moreover, these data resolved disparate reports regarding aneuploid human neurons.

Approaches that measured single (or few) chromosomes in each neuron suggested that >10% of neurons were aneuploid ^72, 73^. Extrapolations based on these data did not account for unmeasured chromosomes in the same neuron, implicitly assuming that every measured aneusomy was unique. We identified 6 aneuploid neurons (2.7%), consistent with other reports ^74, 75^. These and other CNV neurons harbored additional deletions that covered >50% (52/2095) of chromosomes and could be scored by traditional hybridization-based approaches as aneuploid on multiple chromosomes.

In addition to finding a nonrandom distribution of CNVs among CNV neurons, we identified genomic hotspots that were impacted by neuronal CNVs more often than expected by chance; the same approach identified genomic cold spots. Further analysis of these regions found high gene density in cold spots (64.7 +/-56.2 genes per 5Mb region), but a lower gene density (32.6 +/-15.2 genes per 5Mb region) in hotspots. Complementary analysis identified 851 regions with 2 or more CNV breakpoints (*i*.*e*., CNVBs), and found that 220 of these refined previously defined 5Mb hotspots to +/-0.5Mb. Hotspot CNVBs were enriched for long (>100Kb) genes, consistent with the paucity of genes found in these regions. In some cases, the functional consequences of the CNVs are also suggested by associations between long gene expression, neuronal development, and neuropathologies ^76, 77^. For example, we identified seven neurons with distinct CNVs sharing a breakpoint region within *KCNT2*, a long (∼380kb) gene that encodes an outward-rectifying potassium channel. *KCNT2* is important for neuron function and has been linked to several developmental pathologies ^78, 79, 80^ (**Fig. 4B**). *KCNT2* exhibited a TPM of 7.30, which falls within the expected range when considering the expression of all long genes in this tissue (mean TPM 9.56 +/-19.82).

Our study shows that CNV neurons with highly aberrant karyotypes populate neurotypical human frontal cortex. Although their impact on neural circuits and behavior remain unknown, cross-sectional studies indicate that CNV neurons are selectively vulnerable to aging-related loss ^39^.The extent to which recurrent CNV sites are shared among individuals is not yet known; neither is it known if cold sites are refractory to CNV formation or are detrimental to neuronal survival during development. Nevertheless, we report candidate genomic regions that incur frequent neuronal gene rearrangement that provide a rationale for tractable and scalable targeted single cell sequencing. Many interesting questions follow from this study, including whether cold spots in neurotypical individuals are instead aberrant in individuals with neurological disease.

## Acknowledgments

We thank Drs. TE Wilson and FH Gage for essential insight and helpful critique throughout the study, and ML Gage for editorial assistance. We also thank M Wolpert and M Haakenson for technical assistance. Members of the BSMN consortium are listed in a supplementary file, G. Senthil, M Gitik, and T Lehner organized the BSMN consortium. The genome analysis and technology, and flow cytometry cores at the University of Virginia School of Medicine assisted with sample preparation.

## Authors contributions

MJM, REM, JVM, JMK, and DRW designed the study. SBE, IEB, JHS, JMK, and MJM generated sequencing data. CS, KK, BK, JMK, REM, MJM performed data analysis. CS, KK, JVM, JMK, REM, and MJM wrote the manuscript.

## Data and materials availability

Data and call sets have been deposited in the NIMH Data Archive (NDA Study ID 1680, http://dx.doi.org/10.15154/1527774) and can be accessed as part of the NIMH Data Archive permission groups: https://nda.nih.gov/user/dashboard/data_permissions.html. The workflow to generate the final call set is available at https://github.com/mills-lab/Scoval

## Methods

### Sample and sequencing library preparation

We examined human neurons dissected from the dorsolateral prefrontal cortex (DLPFC) of a neurotypical individual (postmortem, 49-year-old male individual, ID: Br5154) used as the common reference brain in a previous study ^28^. Neuronal nuclei (NeuN+) were isolated as in ^39^. We then applied 10X Genomics Chromium Single Cell sequencing that ligated barcodes on the DNA in single cells within a Cell Bead Gel and the barcoded fragments are then pooled for library production, which can profile thousands of cells. We sequenced 2,125 neurons in two batches with mean coverage 0.114X (**Figure 1A**). We further applied 10X Genomics Chromium Linked-Read sequencing to dural fibroblast tissue with very high sequencing coverage (52.7X) from the same individual to identify and phase germline SNPs by isolating and fragmenting long DNA segments into barcoded short reads that could be used to reconstruct underlying haplotypes using Long Ranger v2.2 (https://github.com/10XGenomics/longranger).

### Optimization of Ginkgo for single-cell CNV identification

The final CNV call set was generated using a combination of read-depth and phased loss-of-heterozygosity (LOH)-based validation. First, we processed read alignments from 2,125 single-cells using an adapted version of Ginkgo ^44^ to arrive at our *unvalidated call set*. The call set was then filtered via empirical *P*-value selection using information pertaining to loss of a particular haplotype, obtained by aligning sample reads to the (diploid) phased genome for this individual. The resultant calls were then filtered using a Bayesian classification model to arrive at the final CNV call set, which was further classified by CNV type (heterozygous deletions, homozygous deletions, and duplications) because the strength of support is different for these different CNVs, and the ensuing permutation testing (using heterozygous deletions alone) became more regularized. Only CNV calls in autosomes were included in the final CNV call set. We will now describe the generation pipeline, similar to ^39^, in some detail.

### Setting CNV calling cutoffs in Ginkgo via Gaussian Mixture Model

Ginkgo was optimized by resetting default copy-number cutoffs that determine whether a segment detected by circular binary segmentation (CBS) will be called a CNV. To this end, we processed single cell BAM files from 585 cells obtained from the five control individuals studied in ^39^ using the CBS implementation DNACopy (https://bioconductor.org/packages/release/bioc/html/DNAcopy.html). Aligned reads from each single cell were separately processed into 5,067 autosomal bins across the hg19 human reference genome delineated by Ginkgo, which were then normalized to obtain an average copy number of two for the cell. These individual bins were then grouped contiguously into segments based on similarity of their read coverage using DNACopy. We then fit a Gaussian Mixture Model (GMM) to the distribution of the median copy number of all segments from all cells using an “undoSD” of three, whereby two putative segments had to be more than three times the standard deviation in “intra-segment” copy number to be actually written as separate segments, and alpha=.01. From this fit, the two-tailed probability for the Gaussian curve centered at CN=1 and the one at CN=2 was calculated to be 1.63 (**Supplementary Fig. 1B**). This became the new copy-number cutoff for Ginkgo to call deletions. As seen in **Supplementary Fig. 1B**, there were not many candidate duplications to yield a proper fit, but the duplication cutoff was set at 2.43.

### Filtering to remove outlier bins via Tukey’s rule

Next, the raw bin CN data were filtered for the presence of uniform outlier bins across all cells (*e*.*g*., due to data-specific genomic regions uniformly subject to overamplification or underamplification, regions of poor mappability in the genome, etc). The median of copy numbers of 2,125 cells for each of the 5,067 autosomal bins was first plotted. Tukey’s rule was then applied to tag all bins whose median copy number exceeded Q3 + 1.5* IQR, or was below Q1-1.5*IQR, where the interquartile range IQR ≡ Q3-Q1 and Q1 and Q3 are the first and third quartiles, respectively, of all the median copy numbers. Three hundred and eight outlier bins were identified in addition to Ginkgo’s original list containing 29 (**Supplementary Fig.1C**).

These bins were simply removed from the genome by Ginkgo prior to segment processing while other bins (retaining their genomic coordinates) were merged. For reference, the genomic bin size used by for benchmarking Ginkgo was 500 Kb. Thus, in this work, as in ^39^, we used Ginkgo settings pertaining to an approximate variable bin size of 500 Kb (“variable_500kb_101_bowtie”) and only considered large (> 1 Mb) CNVs. Gingko reported a final mean bin size of 569 Kb, with bins ranging in size from 501 to 2812 Kb.

### Filtering of irregular cells

For all cells, the mean absolute deviation (MAD) of bin copy numbers was calculated and fit to a Gaussian distribution. The mean (mu) and standard deviation (sigma) were .253 and .111, respectively. CNV calls from 19 cells (MAD > mu + 3* sigma) were removed before processing the data further (**Supplementary Fig. 1A**). The total number of reads for all remaining cells ranged uniformly from 580,809 to 8,983,573. However, one cell contained an inordinate proportion of reads (> 80%) aligned to just one of the chromosomes and was removed. Further, eight cells that were not filtered by the above methods were manually curated from the data set based on unlikely copy-number patterns, leaving a total of 2,097 *good neurons* (see **Supplementary Fig. 1D**).

### Assessing the coverage-based single-cell CNV call set

To differentiate between bona fide CNVs and potential false-positives due to coverage fluctuations, we leveraged the long-range haplotype information obtained from the 10X linked-read sequences generated from bulk analysis of matched dural fibroblast tissue. We made use of identified heterozygous SNPs (het-SNPs) and initially segmented the genome using phase blocks of heterozygous SNPs as identified by the linked-read data so that each segment would contain SNPs with consistent haplotype labeling. We then binned these segments further into windows of 20-100 SNPs based on empirical observations of SNP and read coverages. For each window in each cell, we then identified reads that overlapped het-SNPs (herein termed “informative reads”) and noted the allele present on the read. Notably, the coverage in each single cell resulted in a sparse number of informative reads per SNP window, typically resulting in 5-15 reads with specific allele information. Using the inferred haplotype of each overlapped het-SNP, we counted the number of reads present on each of the two haplotypes and calculated the absolute log2 ratio between the read counts if the total number of reads on each haplotype was larger than three. We used this log2 ratio to filter the CNV call set from the previous stage. First, we calculated the median log2 ratio of the windows within the CNV regions in the cells with those CNVs and the median log2 ratio of the windows within the CNV regions but in the cells without those CNVs as a background null model. From these data, we derived an empirical p-value for the observed log2 ratio in the sample with the CNV. We then collated the p-values for each individual CNV to derive a p-value distribution and selected a set of candidate CNVs with a p-value < 0.05.

Next, we randomly permuted 100 sets of “non-CNVs” size-matched to these candidate calls to build a GMM from the underlying median log2 ratios of each CNV/non-CNV region, with the assumption that the two distributions followed two distinct Gaussian distributions. Using the median absolute log2 ratios of the two datasets as the training data, we estimated the parameters of the Gaussians and predicted the posterior probability that the CNV belonged to the CNV distribution using a naive Bayesian classifier. Calls with posterior probability > .99 were selected to process further.

As allele imbalance cannot support the homozygous deletions, we implemented a read-depth ratio measurement to add additional support on the calls. We calculated the read-depth ratio for each bin in every cell based on the bulk sequencing from the same tissue ^28^. The read-depth ratio RDR_b,i_ of bin b and cell i can be calculated as

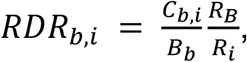

Where C_b,i_ is the number of reads in bin b of cell i, B_b_ is the number of the reads in bin b of bulk sequencing, R_B_ is the total number of reads of bulk sequencing, and R_i_ is the total number of reads of cell i. To distinguish between homozygous and heterozygous deletions, we applied a GMM on read-depth ratio to calculate the posterior probability for the homozygous deletions, and set the cutoff as >0.99 for posterior probability. The final call set for heterozygous deletions was obtained by adjudicating the above calls by requiring the CNV region to have an empirical median log2-ratio p-value (as described above) to be less than .01 (thus ensuring that only calls in regions showing the highest relative allelic preference were selected).

### Benchmarking CNV detection

We applied CHISEL ^47^ to our single-cell sequencing data with its default parameters (max balanced ploidy=4); however, it reported unrealistic results. Only 8.16% of all 5MB windows were reported as normal diploid regions with haplotype copy number ‘1|1’, with most windows (77.83%) indicating the max balanced ploidy with haplotype copy number ‘2|2’. We adjusted the max balanced ploidy setting to 2, resulting in 98.15% of the windows now indicated as normal diploid regions. We combined neighboring CNV windows within the same cell to calculate the overlap percentage with our final call set.

### Clonal cells and recurrent CNVs

To detect the clonal structure of neurons based on CNVs, we designed a very conservative method to identify clonal events. We first found all the CNVs that shared the same start and end breakpoints, then we marked these loci as CNVR. With the haplotype information, we could identify whether these loci were clonal events or the recurrent events that existed on the different haplotypes. For each bin covered by the CNVR, we took the maximum log2 ratio and minimum log2 ratio of the cells with the CNVR and calculated the delta log2 ratio using maximum minus minimum. Next, we calculated the median delta log2 ratio across the bins for each CNVR and observed two distinct distributions, one representing potential clonal events (low delta log2 ratio; CNVs are on the same haplotype) and the other indicating likely independent events (high delta log2 ratio; CNVs are on the different haplotypes).

### Characterizing CNV Neurons

#### Neuronal distribution of CNVs

The raw distribution of the number of CNVs per neuron is shown as a histogram (**Fig. 2A**) on a log scale, along with a null model based on a uniform random distribution of all CNVs in the final call set across all good neurons. Thus, a Poisson curve with mean = (# final CNVs) / (# good cells), scaled up by the total number of good neurons, was superimposed on the first plot to assess whether the final call set contained more CNV-rich neurons than expected by a uniform distribution.

#### Hierarchical clustering and complex karyotypes

The 2,097 good neurons were ordered based on the number of total base pairs affected by heterozygous deletions in descending order. A heat map of all cells was generated showing the percentage of base pairs affected by heterozygous deletions in each autosome (see **Fig. 2C**), Neurons were sorted and numbered in reverse order of % base pairs affected. Those cells affected more than 5% were termed *complex neurons* and numbered 1-65 in our call set. All good neurons were clustered using hierarchical clustering using each autosome as an independent dimension and the percentage of base pairs affected as the distance measure.

Thus, cells with chromosomes that were similarly affected by heterozygous deletions clustered together (**Fig. 2D**). Some cells with possibly multiple recurrent events were identified (**Fig. 2E**), and some seemingly clonal cells were analyzed to be technical replicates.

#### Identifying CNV hotspots and cold spots via permutation testing

The final heterozygous deletion call set was “shuffled” using bedtools ^81^ to arrive at 10,000 unique synthetic permutations (**Fig. 3A**). In each permutation, CNVs in each cell were permuted uniformly at random in the autosomes while prohibiting collision (“noOverlapping” option) and then assembled together. The process was repeated 10,000 times without genomic constraints, as unmappable regions were *a priori* removed (refer to subsection ***Optimization of Ginkgo for single-cell CNV identification***), and calls “straddling” such regions commonly occurred in the final call set (**Supplementary Table 2**).

Each autosome was divided into contiguous 5Mb regions (remaining smaller tails of chromosomes were not considered). The number of unique hits (defined as simple overlap) of each region with synthetic CNVs from all 10,000 permutations was recorded, resulting in a CNV distribution profile for the synthetic data. For each 5Mb region, a *P*-value was assessed for the number of CNV hits in real data among the 10,000 hit-values in the region’s synthetic CNV profile. For our purposes, we define *P*-value to be the fraction of simulated instances that were *at least* as high as the real number of CNV hits to the 5Mb region. Given that CNV hits are discrete-valued, and we are using the same definition of *P*-value for cold spots and hotspots, we impose a more stringent cutoff for cold spots to account for the inherent liberal treatment of data values on the lower extreme (which may lead to an overabundance of cold spots). Regions with a *P*-value < .05 (*i*.*e*., where hits were among the top 5 % of synthetic hit-values for that region) were termed “hotspots” and those with *P*-value > .99 were termed “cold spots.” *Regional significance* (defined as 1 -*P-*value) was plotted against the autosomal genome on the x-axis (**Supplementary Fig. 9**). The distribution of the raw number of CNV hits in 5Mb regions is shown in **Figure 3B**. Cold spots were screened for aberrant genomic blocks that might hamper CNV calling or regions *a priori* neglected. To this end, cold spot regions were coordinate-merged (via “bedtools merge”) and compared to all *a priori* removed bad bins as well as blacklisted regions ^82^ by means of a relative permutation analysis. A merged cold spot that overlapped more with blacklisted regions and bad bins as appropriately compared to 1,000 randomly selected non-cold spot intervals was removed from the list of final cold spots (the cutoff chosen was p > .05) (**Supplementary Fig. 10A)**. Each merged cold spot was mapped to 1,000 randomly selected regions other than existing cold spots, and its overlap with bases contained in bad bins and blacklisted regions, respectively, were calculated in each instance in order to assign it a p-value. For additional relevant detail, some genomic heat maps of copy number of CNV neurons are shown in **Supplementary Figure 10C, D** along with merged cold spots and bad bins. For rigor, cold spots were analyzed for the presence of deduplicated germline structural variants from 1,000 individuals from FusorSV ^83^, the cold spots had a larger SV coverage (11.4) than the unremarkable regions (7.25), further supporting that CNVs are callable in these regions.

Hotspots and cold spots are shown throughout the genome in a Circos ^84^ plot along with 33 regions of the genome where germline CNVs are associated with neurodevelopmental phenotypes ^56^ to assess any possible correlation between the two (**Fig. 3C**). The distribution of the number of genes in 5Mb regions was also plotted for hotspots, cold spots and unremarkable regions as control (**Fig. 3D**). Similar distributions were plotted (with assigned p-values) for long genes and different expression levels (**Supplementary Fig. 11**).

In a complementary assessment, the above permutation analysis was repeated for genes instead of 5Mb genomic regions. To profile gene expression, histograms of p-values for genes were shown for different gene expression categories (**Supplementary Fig. 12)** to assess/confirm general prevalence of hotspots and cold spots in each expression category.

#### Recurrent CNV breakpoint analysis

To assess the impact of different Ginkgo bin sizes on the CNV breakpoint distribution, we used the previously described 10K permuted CNV sets to determine the relationship between the number of breakpoints and Ginkgo bin size. We calculated the mean of the number of breakpoints from all permuted CNVs and compared this to the size of the Ginkgo bin in which they fell. We then normalized the number of breakpoints by the Ginkgo bin size and compared this normalized number of observed breakpoints within CNVB regions with those in permuted regions using a one-sided t-test with the alternative hypothesis that observed > permuted. We then calculated the normalized number of long genes (>100K) overlapped with CNVB bins and compared against the permuted regions using the same strategy. The gene expression analysis was conducted by calculating the transcript per million (TPM) values for the longest gene observed in each of the CNVB and permuted regions and assessing whether they were significantly different using a one-tailed t-test.

