## Supplementary material for "Mapping the Complex Genetic Landscape of Human Neurons": CSun_BioRxiv23_sup.pdf

Chen Sun[1,#] , Kunal Kathuria[2,#], Sarah B Emery[3], ByungJun Kim[1], Ian E. Burbulis[4,5], Joo Heon Shin[2], Brain Somatic Mosaicism Network, Daniel R. Weinberger[2,6,7], John V. Moran[3,8], Jeffrey M. Kidd[1,3], Ryan E. Mills[1,3*], Michael J. McConnell[2*]
[1]Department of Computational Medicine and Bioinformatics, University of Michigan Medical School, 100 Washtenaw Avenue, Ann Arbor, MI 48109, USA.
[2]Lieber Institute for Brain Development, 855 North Wolfe Street, Baltimore, MD 21205, USA.
[3]Department of Human Genetics, University of Michigan Medical School, 1241 East Catherine Street, Ann Arbor, MI 48109, USA.
[4]Department of Biochemistry and Molecular Genetics, University of Virginia, School of Medicine, Charlottesville, VA 22902, USA
[5]Facultad de Medicina y Ciencia, Universidad San Sebastián, Sede de la Patagonia, Puerto Montt, Chile.
[6]Department of Psychiatry and Behavioral Sciences and Neuroscience, Johns Hopkins School of Medicine, 600 North Wolfe Street, Baltimore, MD 21287, USA.
[7]McKusick-Nathans Institute of Genetic Medicine, Johns Hopkins School of Medicine, 733 North Broadway, Baltimore, MD 21230, USA.
[8]Department of Internal Medicine, University of Michigan Medical School, 1500 East Medical Center Drive, Ann Arbor, MI 48109, USA.

### These authors contributed equally
* Communication to

**BSMN Consortium** **1**
**Supplementary Fig. 1** 2
**Supplementary Fig. 2** 4
**Supplementary Fig. 3** 6
**Supplementary Fig. 4** 7
**Supplementary Fig. 5** 8
**Supplementary Fig. 6** 9
**Supplementary Fig. 7** 10
**Supplementary Fig. 8** **12**
**Supplementary Figure 9** 13
**Supplementary Figure 10** 14
**Supplementary Figure 11** 15
**Supplementary Figure 12** 16
**Supplementary Figure 13** 19

**The BSMN consortium:**

**Boston Children's Hospital:** Sara Bizzotto, Michael Coulter, Caroline Dias, Alissa D'Gama, Javier Ganz, Robert Hill, August Yue Huang, Sattar Khoshkhoo, Sonia Kim, Alice Lee, Michael Lodato, Eduardo A. Maury, Michael Miller, Rebeca Borges-Monroy, Rachel Rodin, Christopher A. Walsh, Zinan Zhou

**Harvard University:** Craig Bohrson, Chong Chu, Isidro Cortes-Ciriano, Yanmei Dou, Alon Galor, Doga Gulhan, Minseok Kwon, Joe Luquette, Peter Park, Maxwell Sherman, Vinay Viswanadham

**Icahn School of Medicine at Mount Sinai:** Schahram Akbarian, Andrew Chess, Attila Jones, Chaggai Rosenbluh

**Kennedy Krieger Institute:** Sean Cho, Ben Langmead, Jeremy Thorpe

**Lieber Institute for Brain Development:** Jennifer Erwin, Andrew Jaffe, Michael McConnell, Rujuta Narurkar, Apua Paquola, Jooheon Shin, Richard Straub, Daniel Weinberger

**Mayo Clinic Rochester:** Alexej Abyzov, Taejeong Bae, Yeongjun Jang, Yifan Wang

**Sage Bionetworks:** Cindy Molitor, Mette Peters

**Salk Institute for Biological Studies:** Fred Gage, Sara Linker, Patrick Reed, Meiyan Wang

**Stanford University:** Alexander Urban, Bo Zhou, Xiaowei Zhu, Reenal Pattni

**Universitat Pompeu Fabra:** Aitor Serres Amero, David Juan, Irene Lobon, Tomas Marques-Bonet, Manuel Solis Moruno, Raquel Garcia Perez, Inna Povolotskaya

**University of Barcelona:** Eduardo Soriano

**University of California, Los Angeles:** Gary Mathern

**University of California, San Diego:** Danny Antaki, Dan Averbuj, Laurel Ball, Martin Breuss, Eric Courchesne, Joseph Gleeson, Xiaoxu Yang, Changuk Chung

**University of Michigan:** Sarah B. Emery, Diane A. Flasch, Jeffrey M. Kidd, Huira C. Kopera, Kenneth Y. Kwan, Ryan E. Mills, John B. Moldovan, John V. Moran, Chen Sun, Xuefang Zhao, Weichen Zhou, Trenton J. Frisbie, Yifan Wang

**Yale University:** Adriana Cherskov, Liana Fasching, Alexandre Jourdon, Sirisha Pochareddy, Soraya Scuderi, Nenad Sestan, Flora M. Vaccarino

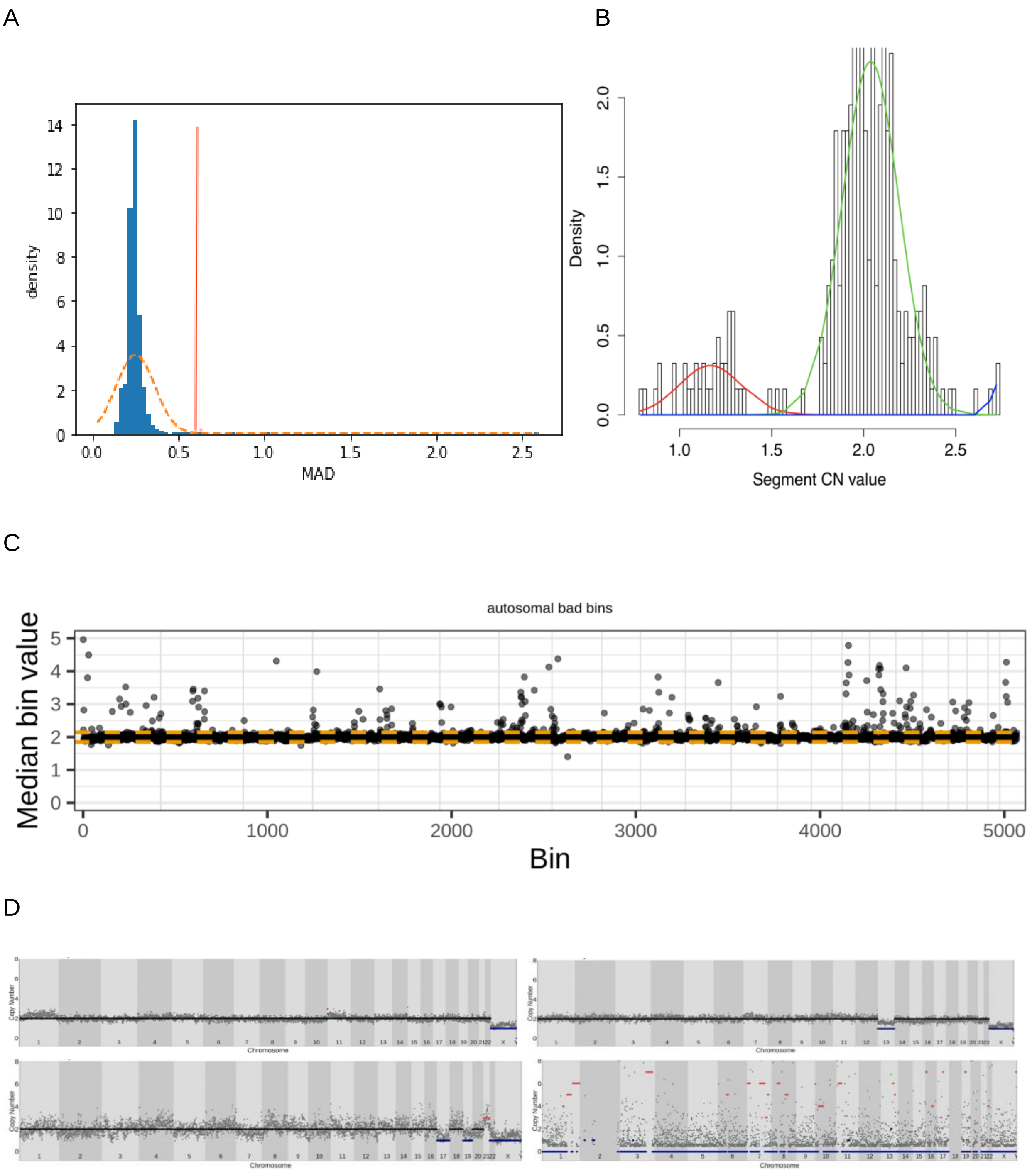


Supplementary Fig. 1

**Optimization of Ginkgo for read-depth-based CNV calls** (**A**) Mean absolute deviation (MAD) score distribution (based on bin copy numbers) excluded 19 of 2,125 neurons (with MAD > 3 standard deviations away from mean). (**B**) Thresholds for calling putative CNVs were set using a GMM based on 585 cells obtained from the 5 control individuals studied in (*20*) at 1.63 for deletions and 2.43 for duplications. (**C**) Tukey's rule was applied to median copy numbers for all genomic bins across all neurons in our dataset to yield 308 additional outlier bins in addition to Ginkgo's original 29 that were excluded from further analysis. (**D**) 4 additional cells that passed the MAD cutoff but were curated manually due to unlikely copy-number patterns, including 1 that did not pass the read-count filter due to concentration of reads on Chr2 (bottom right)

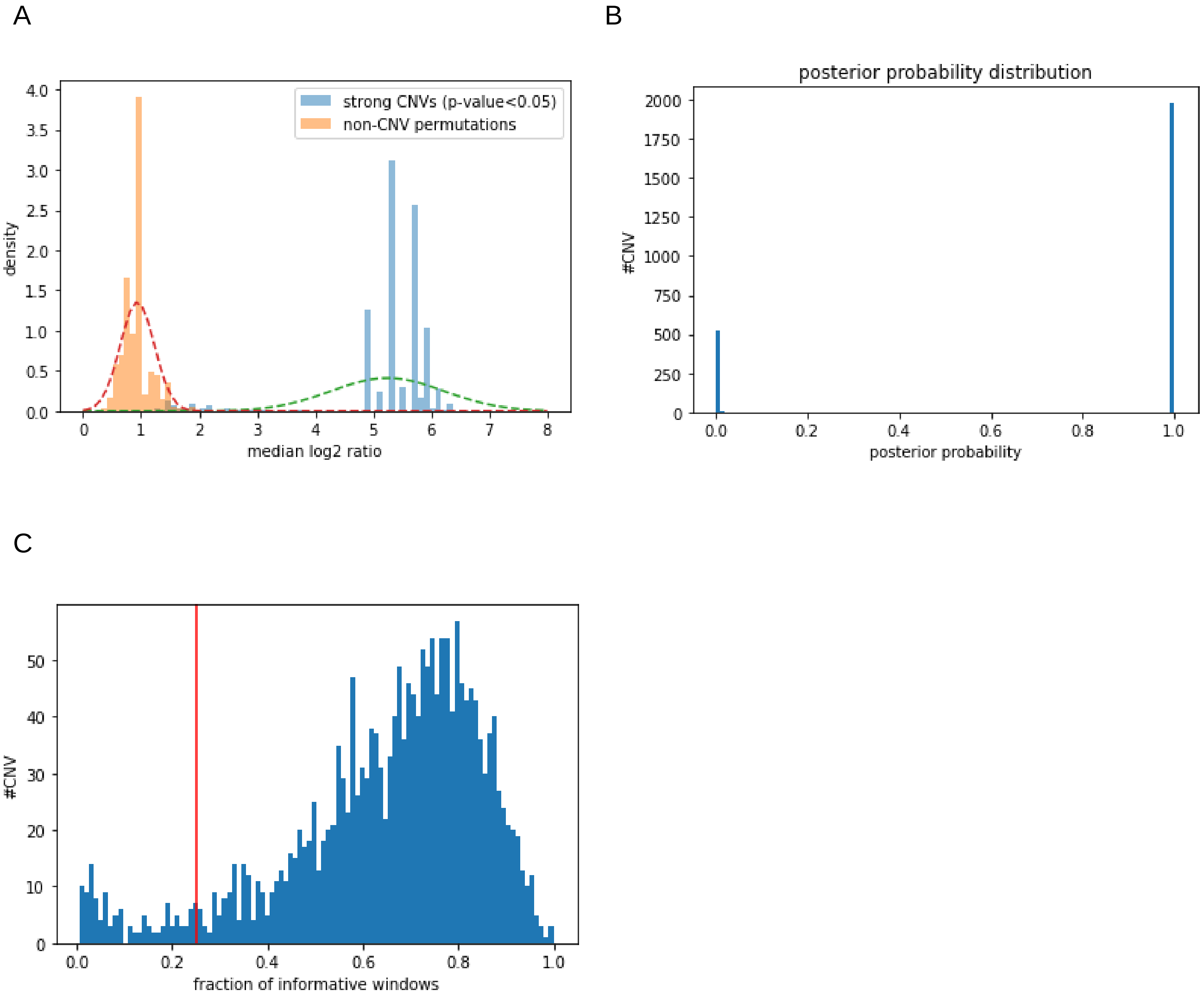


#### Supplementary Fig. 2

**Naïve Bayesian-based pipeline to filter CNVs.** (**A**) We labeled CNV calls as “strong” based on an empirical p-value, which is derived from the median absolute log2 ratio of the windows within the CNV regions. Then we derived a Gaussian mixture model of strong calls and 100 non-CNV set permutations. (**B**) Using the median absolute log2 ratios of the two datasets as the training data, we estimated the parameters of the Gaussians and predicted the posterior probability that a candidate CNV belonged to a specific CNV distribution. (**C**) We filter out deletion calls where more than 75% of its het-SNP windows contained fewer than 3 informative reads which precludes an accurate haplotype assessment.

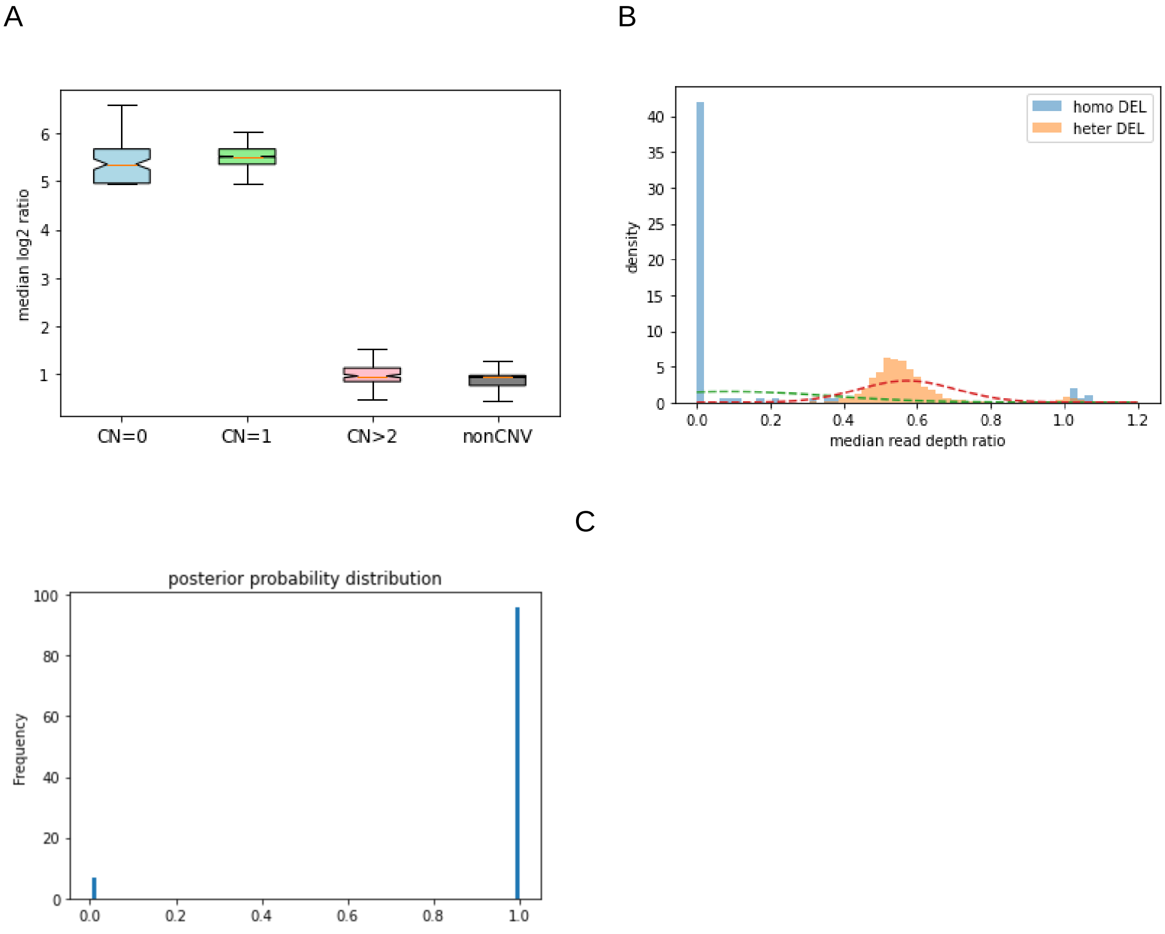


#### Supplementary Fig. 3

**Homozygous deletions and duplications are more challenging to validate using SCOVAL.** (A) The median absolute log2 ratio of informative reads in candidate homozygous deletions informative reads are similar to heterozygous deletions. The median absolute log2 ratio of duplications are not significantly different from randomly sampled non-CNV regions. (B) Derived Gaussian mixture model from median read depth ratios between homozygous and heterozygous deletions. (C) Posterior probability for putative homozygous deletions using a naive Bayesian classifier on the Gaussian mixture model from the initial heterozygous and homozygous deletion calls.

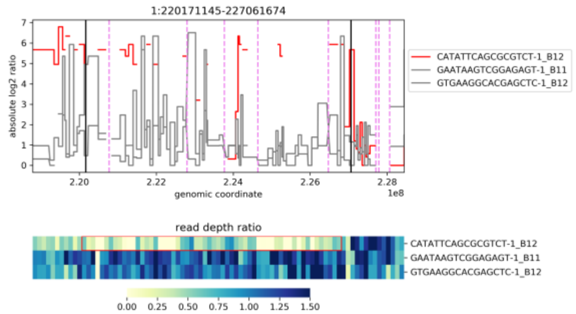
1:220171145-227061674
absolute log2 ratio
genomic coordinate
CATATTCAGCGCGTCT-1_B12
GAATAAGTCGGAGAGT-1_B11
GTGAAGGCACGAGCTC-1_B12
read depth ratio

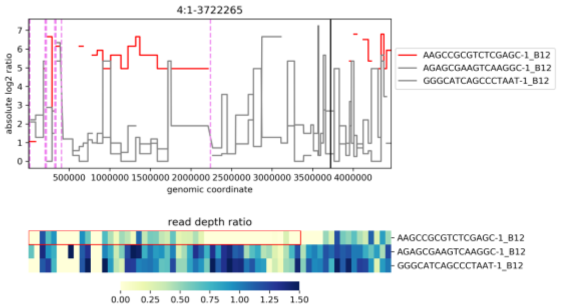
4:1-3722265
absolute log2 ratio
genomic coordinate
AAGCCGCGTCTCGAGC-1_B12
AGAGCGAAGTCAAGGC-1_B12
GGGCATCAGCCCTAAT-1_B12
read depth ratio

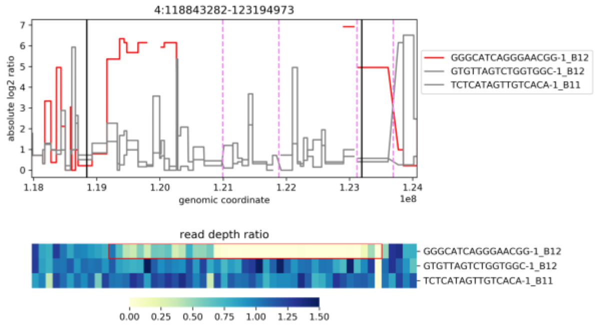
4:118843282-123194973
absolute log2 ratio
genomic coordinate
GGGCATCAGGGAACGG-1_B12
GTGTTAGTCTGGTGGC-1_B12
TCTCATAGTTGTCACA-1_B11
read depth ratio

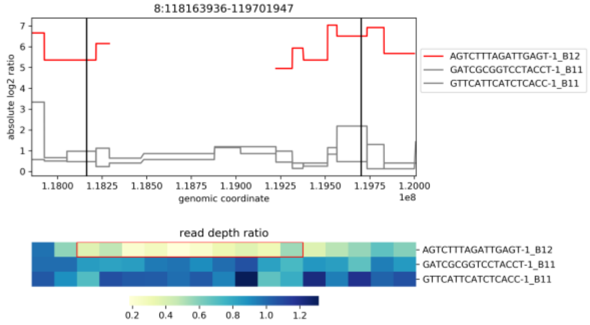
8:118163936-119701947
absolute log2 ratio
genomic coordinate
AGTCTTTAGATTGAGT-1_B12
GATCGCGGTCCTACCT-1_B11
GTTCATTCATCTCACC-1_B11
read depth ratio

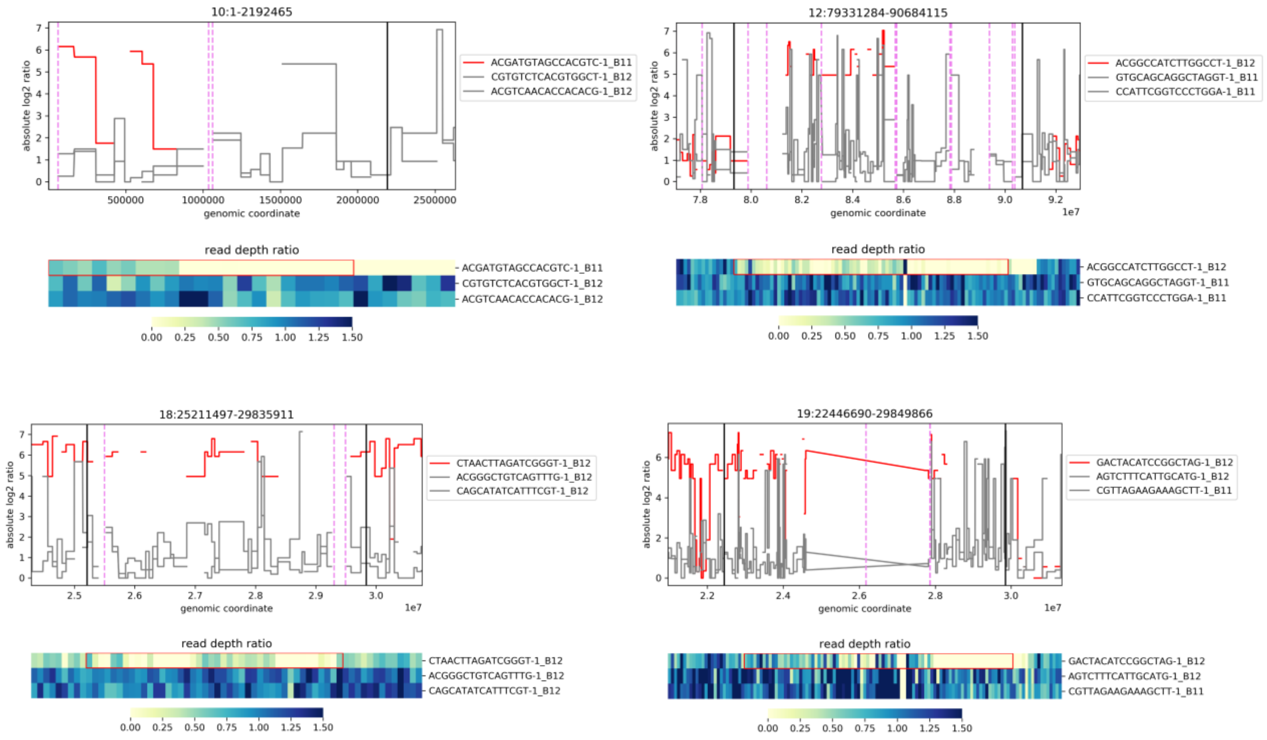


Supplementary Fig. 4

**Heterozygous deletions miscalled as homozygous deletions.** We identified 8 homozygous deletion calls from Ginkgo with read depth and allele ratio characteristics consistent with heterozygous deletions. The upper panel for each figure is the absolute log2 ratio. Red line indicates the cell with CNV and the gray lines represent two random background cells. The bottom panel is the read depth ratio. The first row is for the cell with the candidate CNV, supplemented in rows two and three with randomly chosen cells as background.

A

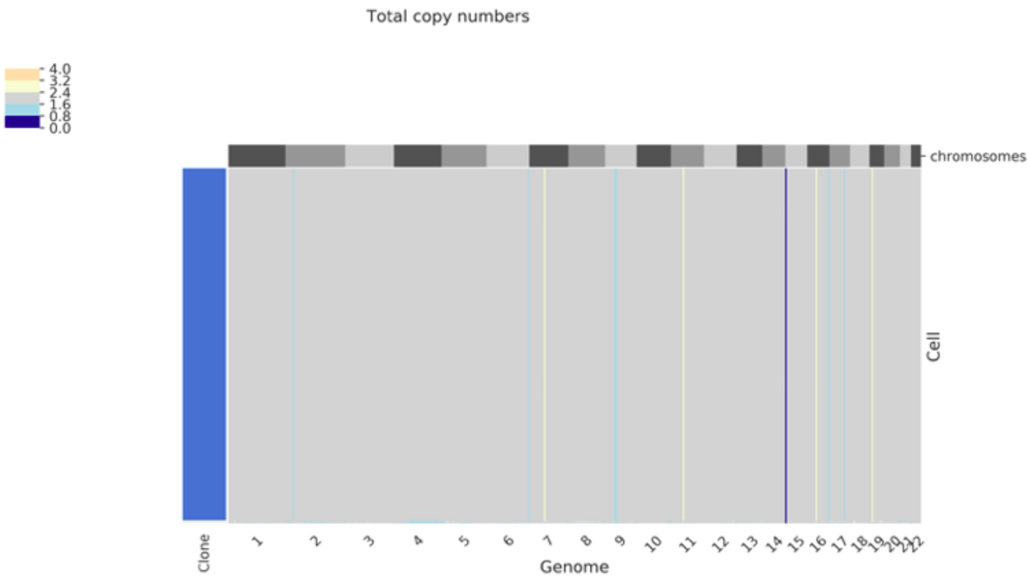


B

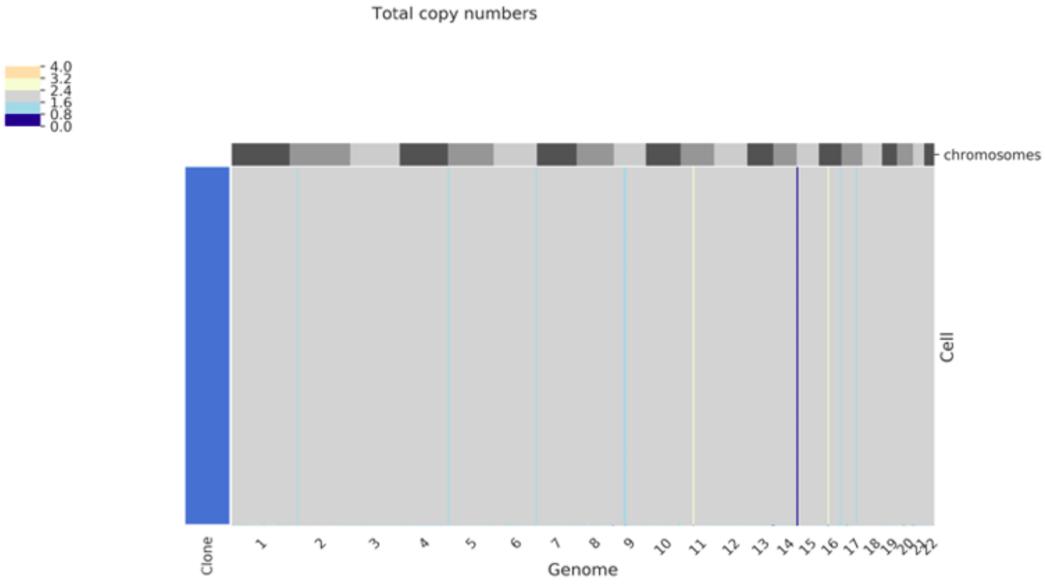


#### Supplementary Fig. 5

**Benchmarking CNV detection with CHISEL.** Output of CHISEL to our single-cell sequencing data in (**A**) batch B11 and (**B**) batch B12 using diploid=2 parameters. The majority of CNVs were reported in all cells.

A

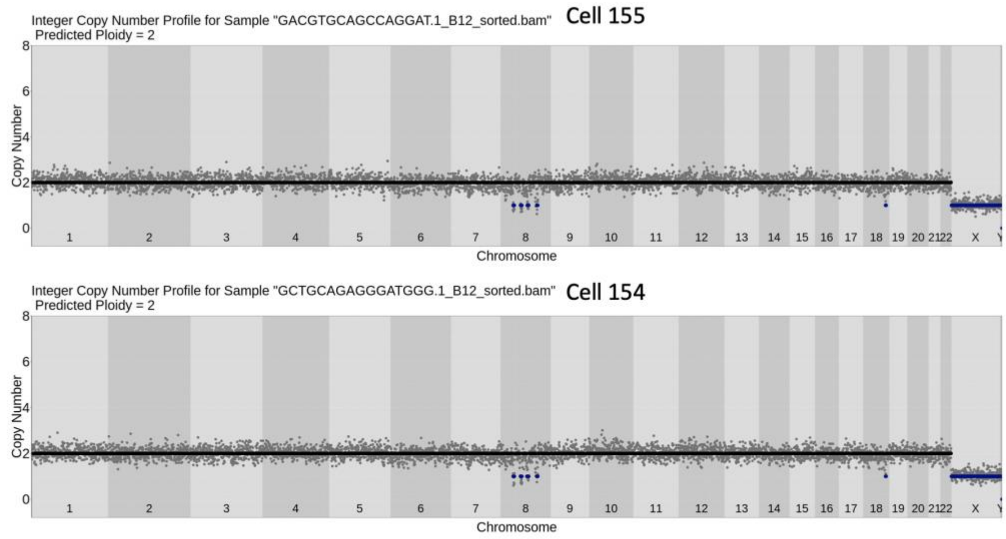


B

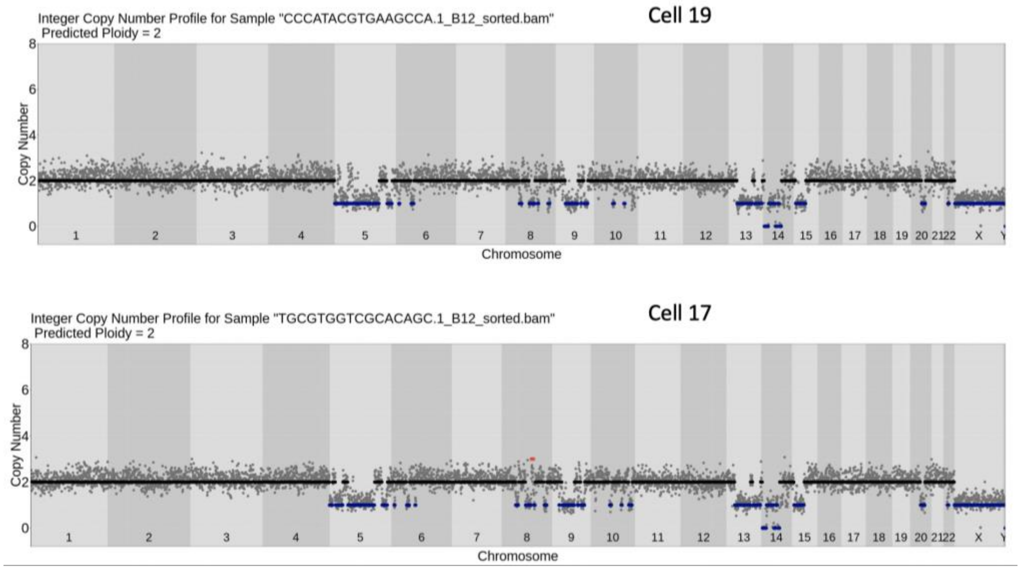


#### Supplementary Fig. 6

**Putative clones that cannot be ruled out as technical replicates**. (**A**) Putative clone-pair 1 (**B**) Putative clone-pair 2. Cells #17 and #19 show some deviances in overall bin copy number variances, but cannot conclusively be established as independently amplified neurons

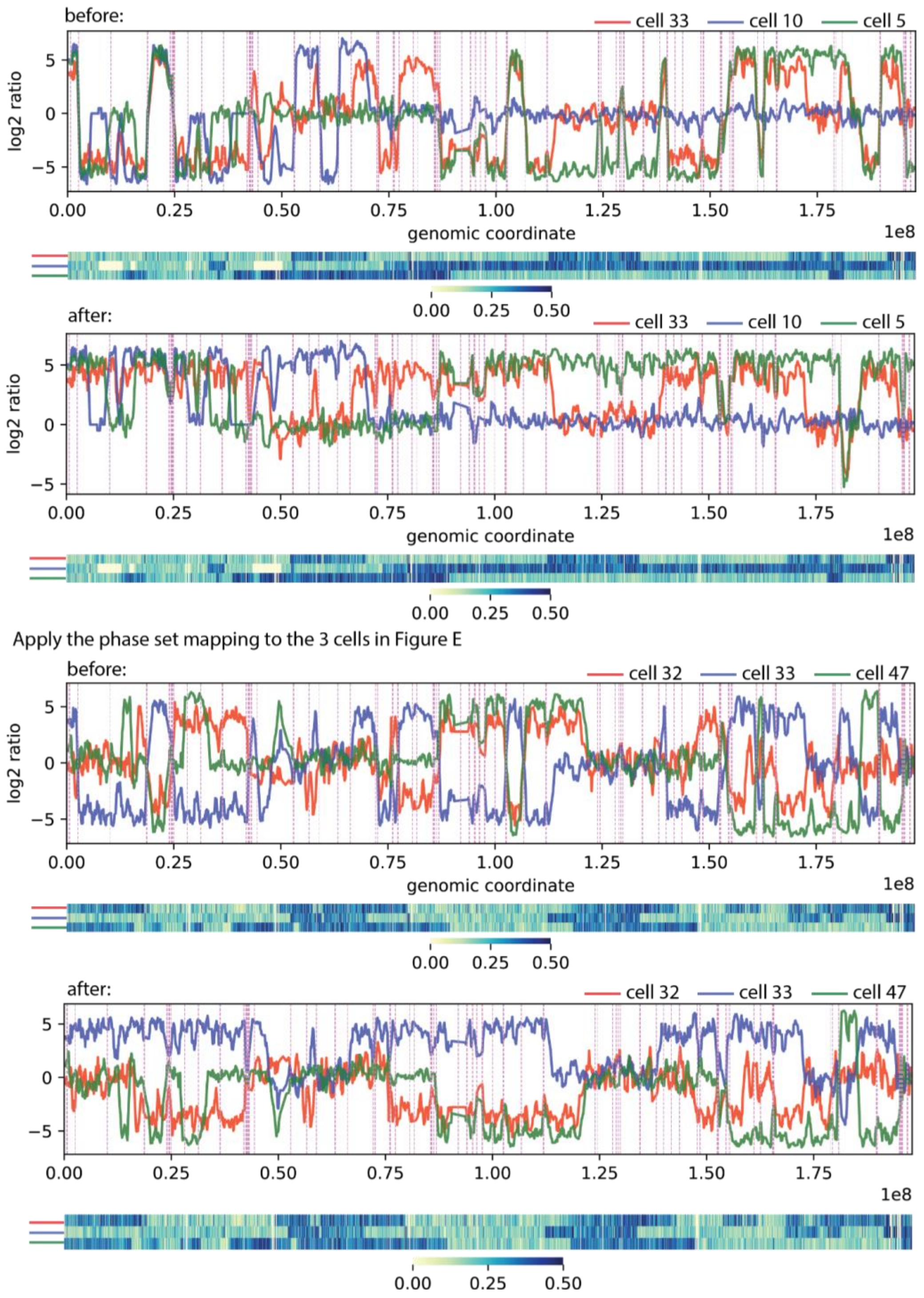


Supplementary Fig. 7

**Reconstruction of Chromosome 3 haplotypes using overlapping heterozygous deletions in 3 cells.** We generated extended phase blocks using three CNV neurons (cells #33, #10, and #5) that contained overlapping deletions that in aggregate cover the full-length of Chromosome

3 in order to determine phasing at chromosome level. These were used to reconstruct the haplotype of 3 cells reported in Figure 2E.

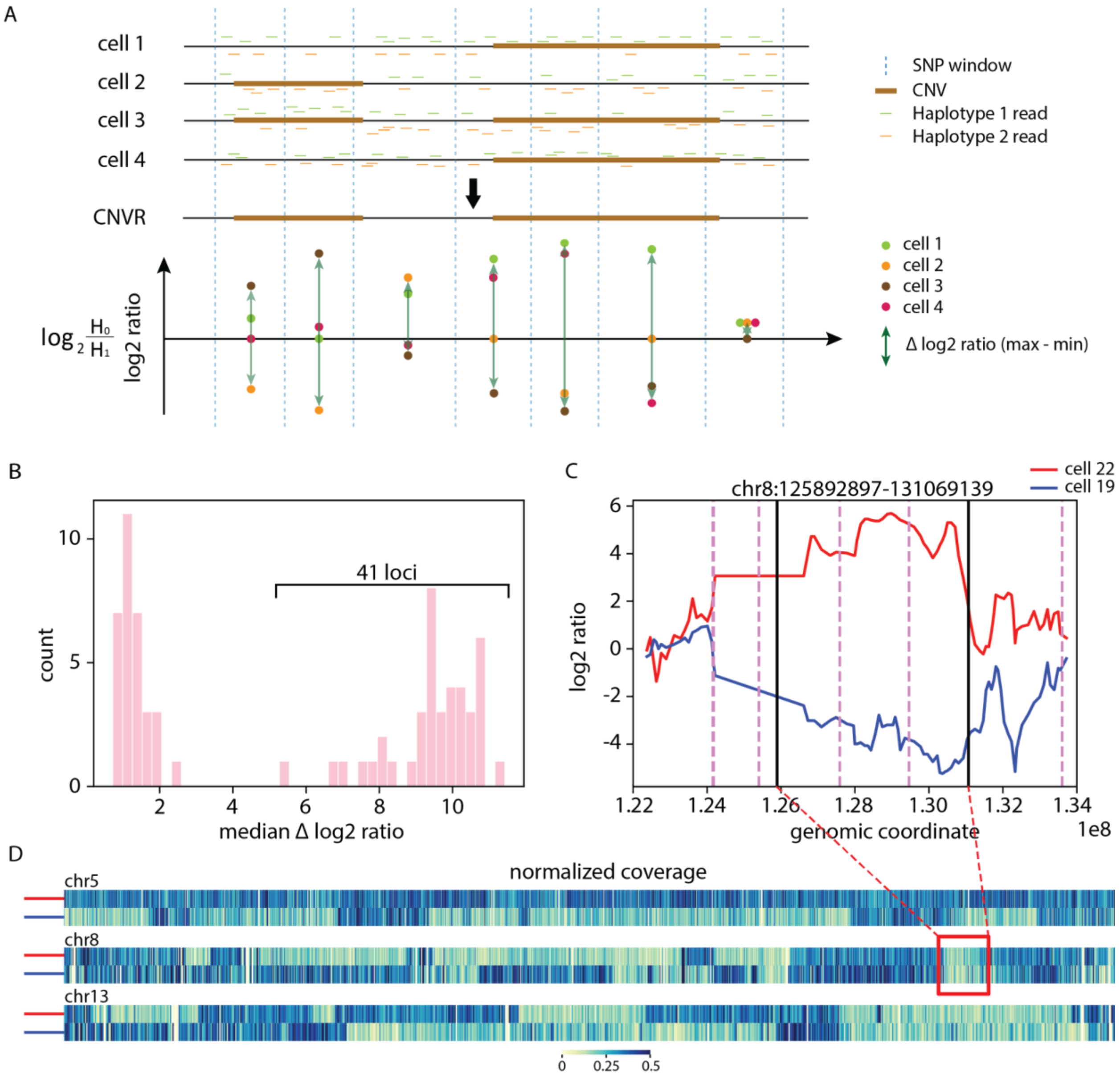


Supplementary Fig. 8

**CNVs sharing the same location are on different haplotypes**. (**A**) We derived a min-max median delta log2 ratio to determine whether CNVRs likely reside on the same haplotype. (**B**) There are two apparent distributions of delta log2 ratio values. CNVs from 41 CNVRs with higher median delta log2 ratio likely occurred on different haplotypes. (**C**) Two cells (#22 and #19) both exhibit CNVs with the same location on Chr8, but show allelic ratios consistent with residing on different haplotypes. (**D**) An examination of CNVs on other chromosomes in these cells further indicate that these shared CNVs are not clonally derived.

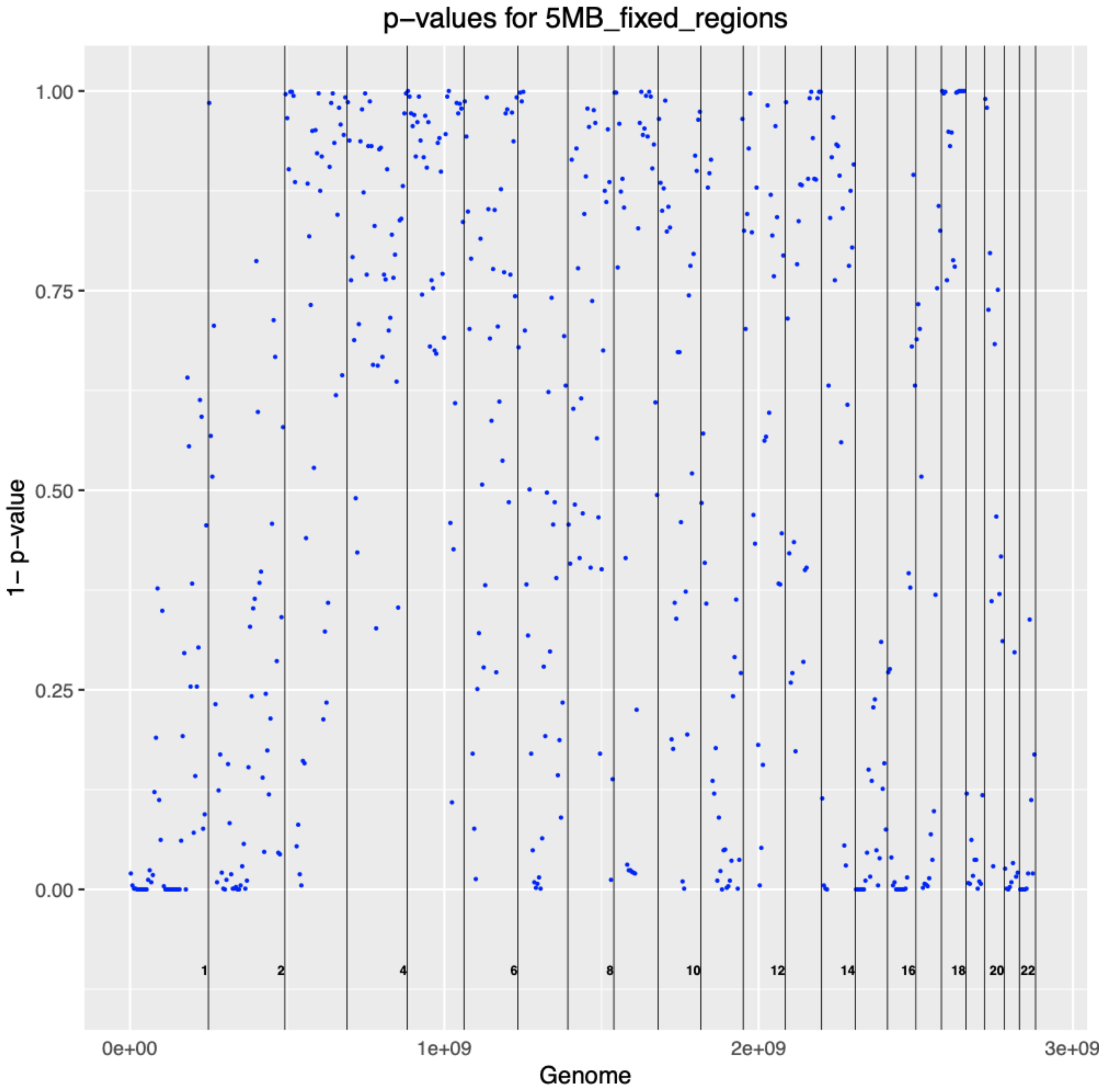


#### Supplementary Figure 9

**Genome-wide identification of hotspots and cold spots**. Regional significance (1 minus p-value of number of CNV hits in a 5Mb region compared to random synthetic data) is plotted for all 5Mb genomic regions. Hotspots and cold spots shown in Fig. 3C. were identified using values > .95 and < .01 respectively.

A

Overlap of bad bins with cold spot permutations

B

Overlap of blacklisted regions with cold spot permutations

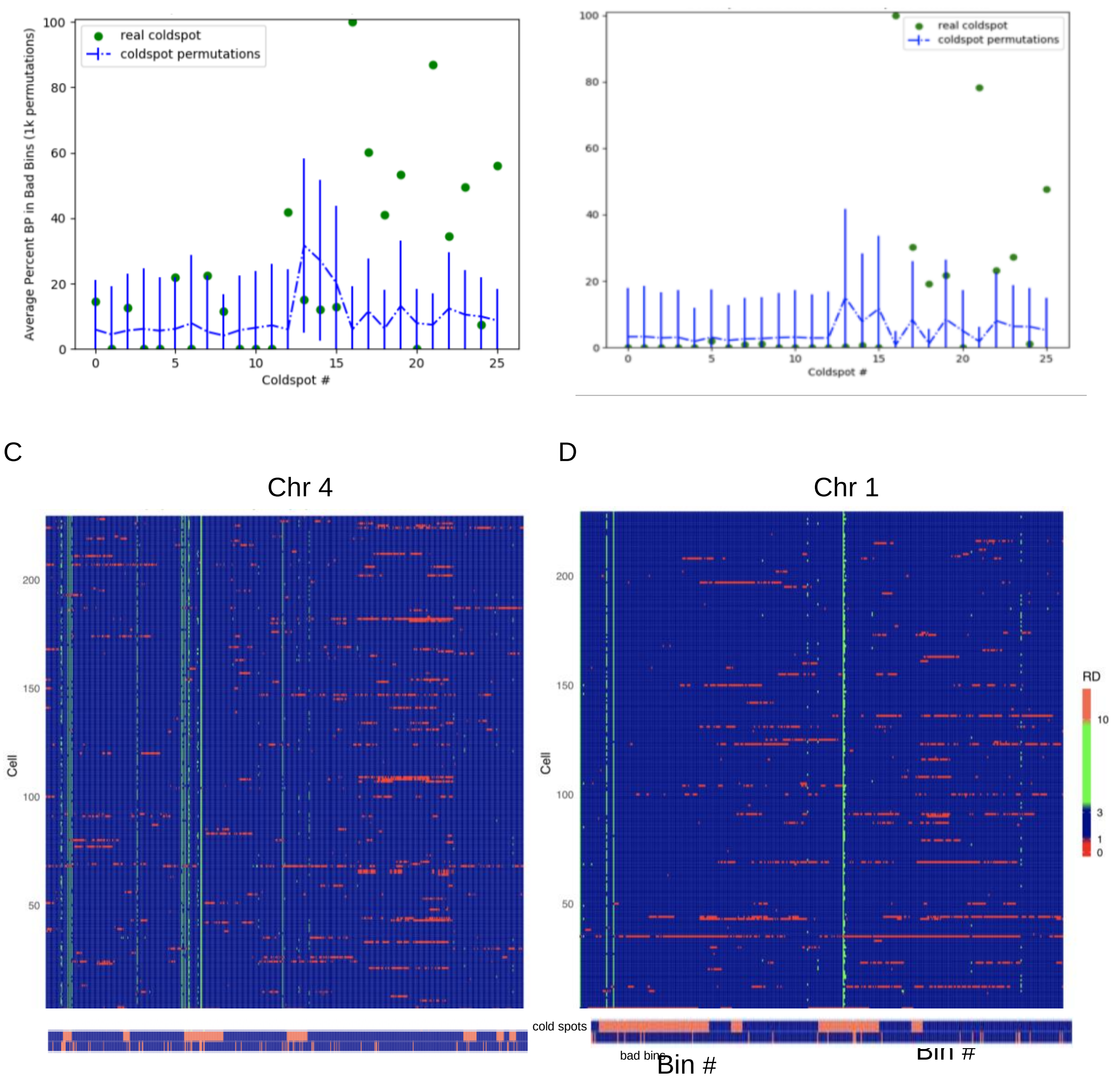


#### Supplementary Figure 10

**Filtering of cold spots based on unmappable genomic regions** (**A,B**) Percentage of cold spots occupied by bad bins and by blacklisted regions identified by ENCODE respectively (green) compared to median of same quantity for cold spot permutations in control regions (blue). Cold spots registering high unmappable content (p-value < .05 cutoff) were filtered out (see **Methods**). (**C,D**) Schematic overview showing correlation of cold spots, bad bins and read depth for all CNV neurons across all genomic bins for 2 different chromosomes.

A

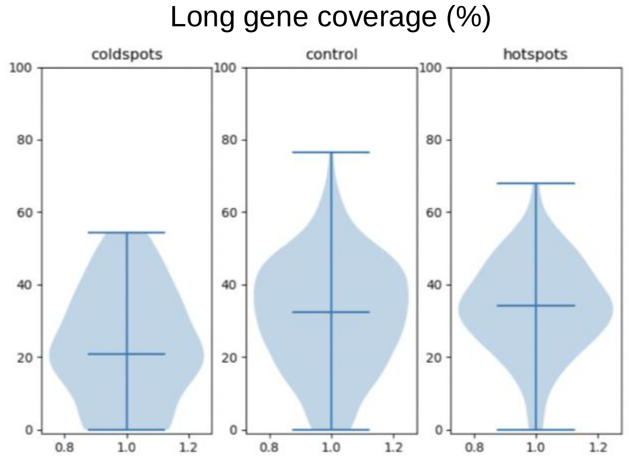


B

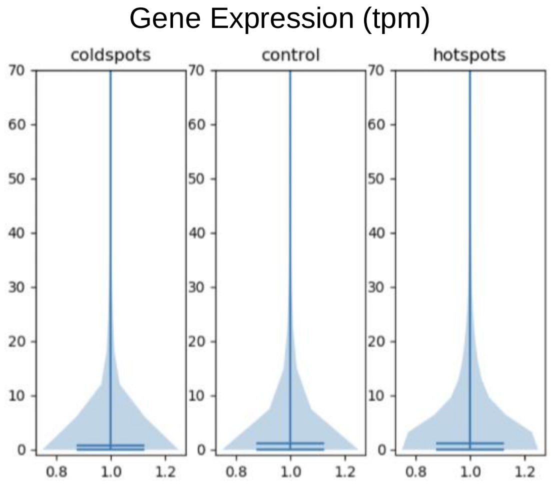


#### Supplementary Figure 11

**Comparison of hotspots and coldspots to control regions regarding long gene coverage and gene expression level**. Violin plot distribution showing (**A**) a depletion of long genes (> 100 Kb) covering the region and (**A**) overall gene expression of genes in the region relative to control (middle) and hotspots (right).

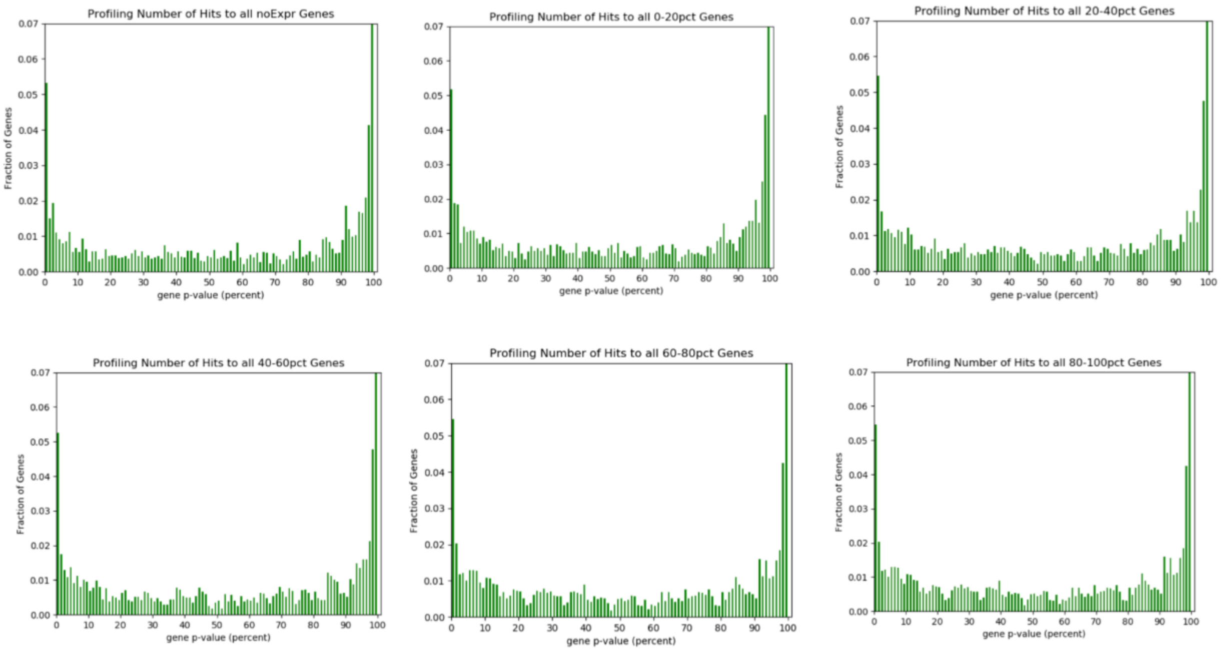


#### Supplementary Figure 12

**Complementary view of hotspots and cold spots in physical genes**. Depicted are the *distribution* of p-values (defined as in Supplementary Figure 9 but for physical genes) for genes showing the presence of hotspots (first 5 bins) and cold spots (last bin) in six expression categories (genes not expressed, and 5 quintiles of genes expressed in DLPFC). This analysis is complementary to that performed in 5Mb regions and shows the presence of hotspots and cold spots in genes expressed at various levels

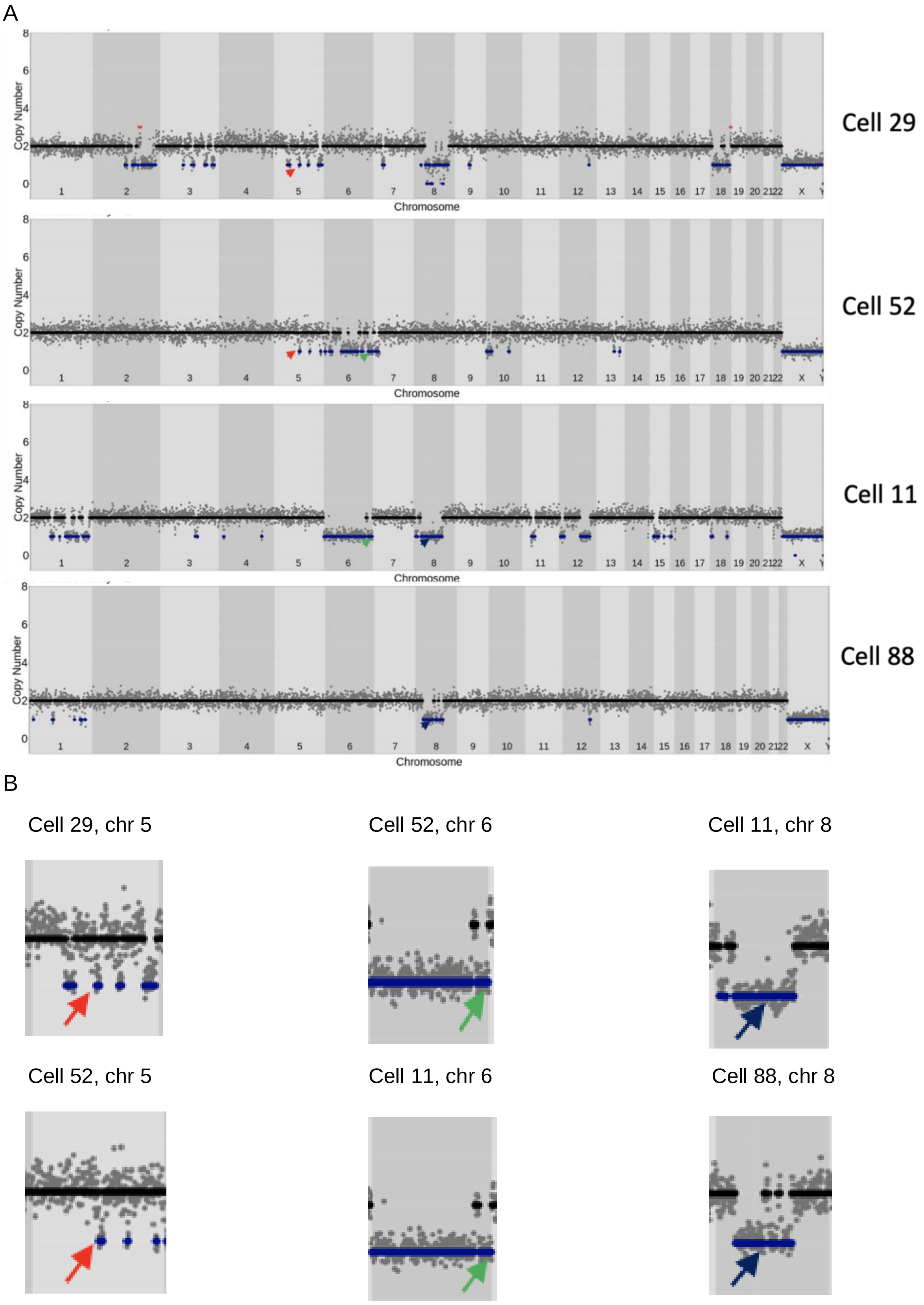
A
Cell 29
Cell 52
Cell 11
Cell 88
Copy Number
Chromosome
B
Cell 29, chr 5
Cell 52, chr 6
Cell 11, chr 8
Cell 52, chr 5
Cell 11, chr 6
Cell 88, chr 8

C

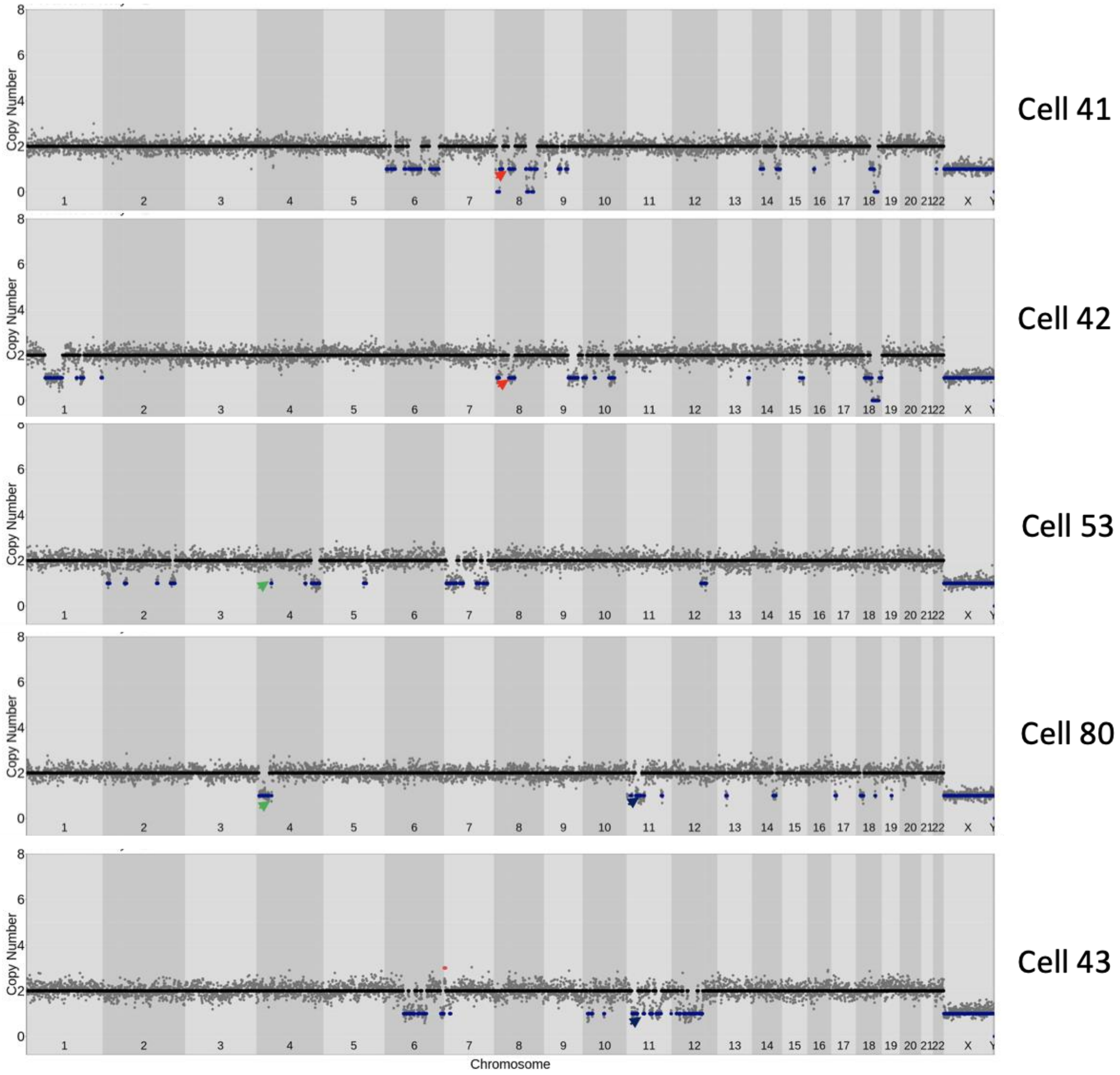
Copy Number
Chromosome
Cell 41
Cell 42
Cell 53
Cell 80
Cell 43

D

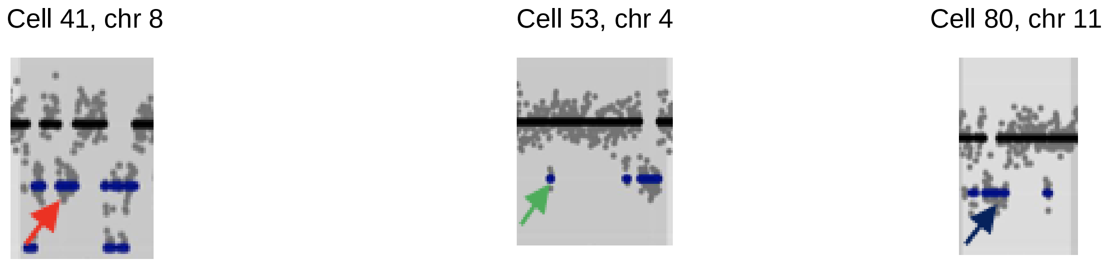
Cell 41, chr 8
Cell 53, chr 4
Cell 80, chr 11

Cell 41, chr 8 Cell 80, chr 4 Cell 43, chr 11

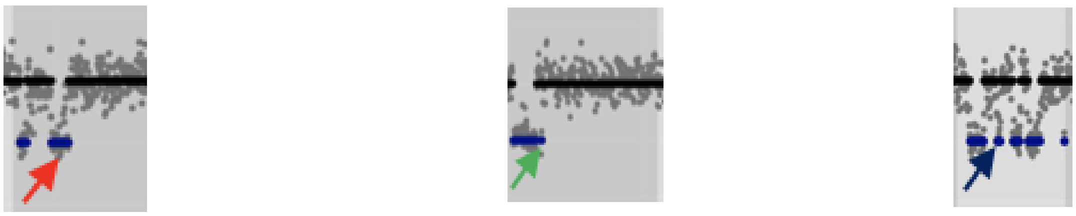

#### Supplementary Figure 13

**Cells showing single shared events among complex karyotypes**. (**A,B**) Three pairs of CNVs in 4 cells (shown by red, green and blue arrows respectively) are shared/recurrent. The shared CNVs are magnified in the lower panel. None of the other CNVs are shared. This indicates that the recurring CNVs are not necessarily clonal. (**C**) Same as above for another group of 5 cells. (**D**) The lower panel shows a magnified view of shared CNVs in (C)
